# Breast cancer spheroids prefer activated macrophages as an accomplice: An in vitro study

**DOI:** 10.1101/2024.09.24.614655

**Authors:** Abhishek Teli, Ranjani Iyer, Karthik Shanbhag, Rahul Gawarguru, Sukanya Gayan, Sumaiya Shaikh, Anup Tamhankar, Siddhesh S. Kamat, Tuli Dey

## Abstract

Cancer, a heterogeneous disease in nature, often requires help from diverse pro-tumor or tumor-associated- cells, which are recruited and persevered within the stroma. Pro-tumor stromal cells provide the essential support needed for tumor growth, metastasis, and development of drug resistance in due time. Tumor-associated macrophages, one of such cells, are essential to tumor microenvironment and tumor survival. In recent years, TAMs have been identified as potential drug targets and therapeutic agents, which encourages the in-depth characterization of their crosstalk with the tumors. The current study has successfully developed a cost-effective in vitro platform for Chemokine Assisted Recruitment of Macrophages to spheroids mimicking the physiology of TAM recruitment. Firstly, monocytic cell line (U937) were converted into activated naive macrophages (M0) and pro-and anti-inflammatory (M1 and M2) subtypes. Monocytes, M0, M1, and M2 macrophages are characterized extensively. Secondly, the naive and polarized macrophages were subjected to chemokine-dependent recruitment into monotypic and heterotypic breast cancer spheroids. The nature of the recruitment is further investigated by assessing the profile of chemokines and chemokine receptors. Recruited macrophages are also observed to manipulate spheroid behavior in many ways. The recruited macrophages also exhibit an increased level of Siglec-1 (CD169), one of the potential TAM markers. The current platform’s potential for application can be extended to understand the recruitment process of other immune/stromal cells to solid tumors. It could be a potential addition to the arrays of in vitro platforms developed to screen the efficiency of cell-based immunotherapeutics in the future.

## Introduction

Cancer, one of the major non-communicable diseases, has the highest death toll to date, which is further predicted to increase significantly by 2050. As per recent estimations, the incidence rate has risen to one in five men or women in their lifetime, while the death rates are about one in nine men and one in twelve women (Bray et al., 2024). Among all kinds of cancers, breast cancer dominates the mortality rate among women, not only in developed countries but also in developing countries like India (Mehrotra and Yadav, 2022). In recent studies, the trend of genetically transmittable cancer (breast) is observed to be increased compared to the infectious ones (cervical) among women (Sung et al., 2021). To date, the majority of solid tumors, including sarcomas and carcinomas, can only be successfully treated with surgery and chemotherapy. However, several factors, including the increasing cost of treatment (Pisu et al., 2018), side effects, developing drug resistance, and chance of relapse, have obstructed the conventional therapeutic approaches (Anand et al., 2023).

In the last decade, small ligand-based and cell-based immunotherapy has been hailed as the ‘breakthrough’ advancement over chemo- and other existing therapies (Hamdan and Cerullo, 2023). In spite of high expectations, the results remain lackluster (Sharma et al., 2017). Among the most anticipated immunotherapeutic modalities, namely the CAR-T therapy, which works against specific subsets of leukemia or lymphoma, seems to give rise to CD19-negative relapse (Aparicio-Pérez et al., 2023). Failure of CAR-T cell-based immunotherapeutics against solid tumors can be attributed to multiple factors, including minimal infiltration, off-target toxicity (Sterner & Sterner, 2021), T cell-mediated cytotoxicity (Schubert et al., 2021), along with the local immune-suppressive TME (Maalej et al., 2023). In this context, other members of the innate immunity system, such as macrophages and Natural killer cells, are coming up as potential candidates for cell-based immunotherapy (Liang et al., 2023).

Among them, macrophages have been identified as an integral part of the tumor immune microenvironment (TIME) and characterized as ‘Tumor-associated macrophages’ or TAMs (Mantovani et al., 1992). Regardless of its innate immunogenic role and phagocytic ability, TAMs are found to perform many pro-tumor activities and support the immune-suppressive environment. Many researchers are trying to understand the potential of TAMs both as therapeutic agents and as targets. Activation of anti-tumorigenic macrophages (M1 type) or abolishment of pro-tumorigenic macrophages (M2 type) have been utilized to evaluate their anti-cancer potential (Mantovani et al., 2022). Macrophage-mediated inflammation and its role in tumor aggression have also become an interesting point of study in recent years (Blériot et al., 2024).

In spite of many pivotal studies on tumor-associated macrophages and their supportive role in tumor behavior, including metastasis, angiogenesis, drug resistance, and others (Pan et al., 2021), many questions remain unanswered. Though the tissue-resident macrophages and TAMs are known to derive from circulatory monocytes, the exact molecular mechanism behind the conversion has yet to be discovered. For tumor-specific recruitment, the CCL2-CCR2 axis was presumed to play an important role (Qian et al., 2011). However, the failure of many CCL2 inhibitors (CNTO888, MLN1202) at clinical trials highlighted the gap in knowledge. In a recent study, Cassetta et al., projected the role of CCL8 as a potential recruiter (Cassetta et al., 2019) which remains to be tested further. Furthermore, the accurate identity of surface markers of tumor-associated macrophages, along with the ‘conventional’ M1/M2 signatures, are also being questioned by several recent studies (Cassetta et al., 2019; Singhal et al., 2019). Few recent studies have reported the preferential recruitment of activated macrophages and tumor-associated monocytes by the tumor (Zhang et al., 2022; Adebowale et al., 2023; Du et al., 2024).

As it happens, most of the TAM-related in vitro studies are done using two-dimensional cultures of cancer cells along with terminally differentiated monocytes (either M1 or M2). Preliminary studies used already differentiated macrophages (Engstrom et al., 2014) or cancer cell-derived media-treated monocytes (Hagemann et al., 2006; Sousa et al., 2015; Benner et al., 2019), which left no opportunity for cancer cell-macrophage crosstalk and cancer cell-mediated differentiation of macrophages. Utilization of 3D models for tumor and macrophage interaction started early (Hauptmann et al., 1993; Konnur et al., 1996) without any prolific development. The development of microfluidics facilitates the direct-co-culture method for multiple cell types; however, resulted in the development of contradictory data (Hsu et al., 2012; Li et al., 2017). For in vivo studies, Trim24 KO mice (Liu et al., 2017), athymic (Zhou et al., 2019), and immunocompetent mouse models (Yu et al., 2019) have been reported earlier. However, the outcome of these studies remains inconclusive as many identified potential therapeutic targets failed in human trials, probably either due to the over-dependence on existing mouse models (Gengenbacher et al., 2017; Olson et al., 2018) or to the lack of an in vitro human-relevant tumor-macrophage crosstalk model (Dey, 2020).

The current project is designed to address the question of developing an in vitro cost-effective human-relevant tumor-macrophage interactive model using breast cancer spheroids (Gayan et al., 2017, 2021, 2024). The spheroids are employed to recruit monocytes and monocyte-derived macrophages, followed by their conversion into TAMs. Moreover, the developed platform will be used to analyze tumor-macrophage interaction in the context of influencing spheroid biology and behavior.

## Result and Discussion

Growing tumors actively recruit circulating monocytes or monocyte-derived macrophages from the circulatory system. Available in vitro models majorly utilized already polarized M2 (anti-inflammatory) subtypes for enhanced incorporation into the tumor or organoid (Dwyer et al., 2016; Hacker et al., 2023; Mazan and Marusiak, 2024). However, as suggested by Cassetta et al., tumor cells presumably start the recruitment process early with the conversion of circulatory monocytes of both classical (CD14++/CD16+) or non-classical (CD14+/CD16+) origin (Cassetta et al., 2019). It is yet remains to be clear, whether the conversion of recruited monocyte-derived macrophages to Tumor-associated Macrophages happens during the journey or after their entrapment within the tumor. A human-relevant in-vitro tumor-macrophage interaction platform is established to assess the tumor-driven recruitment of monocytes and different subtypes of monocyte-derived macrophages (M0, M1, and M2). Here we have utilized monotypic (MCF7 or MDA MB231) and heterotypic (MCF7 and MDA MB231) spheroids from adenocarcinoma cell lines (SI Fig 1) to represent the solid breast tumor physiology.

As observed from the literature, the majority of the tumor-macrophage studies utilized macrophages developed from either THP1 or U937 cell lines. Though both of them are used ubiquitously to generate macrophages, recent studies highlighted that the THP1 generates a pro-M1 population post-activation, whereas U937 cells produce a pro-M2 population of macrophages (Nascimento et al., 2022). We used the U937 cell line for the current study due to its mature nature, originating from an older source and having a predisposition towards the M2 subpopulation as pro-tumorigenic monocytes presumably perform better (Zhang et al., 2022). Additionally, these cells reportedly belong to classical origin (CD14+/CD16-) (Szittner et al., 2013), which is the major source of human TAMs (Olingy et al., 2019). Different sub-types of macrophages have been developed from the human pre-monocyte cell line (U937) following the previously described protocol with minor modifications (Lund et al., 2016; Kuno et al., 2020). The activated and polarized macrophages were characterized to ascertain their subtypes.

### Differentiation and characterization of pro-and anti-inflammatory subtypes of macrophages

As observed in Fig 1A, the activated (M0) and differentiated macrophages (M1 and M2) exhibit multi-nucleation and ruffling of cell membranes compared to the mononuclear monocytes. The increment of cell size in both pro- and anti-inflammatory (M1 & M2) subtypes are also apparent from the images. Ruffling of the actin-rich membrane into macropinosome (fried egg appearance) further highlights the active status of the differentiated macrophages (Quinn et al., 2021). Increased size and granularity of differentiated macrophages (M1 and M2) are observed compared to monocytes and M0 macrophages (Fig 1B), as reported earlier (Buchacher et al., 2015). However, the differentiation efficiency is observed to be different among different batches and never reached 100%, a trend similar to earlier studies (Nascimento et al., 2022).

**Figure 1:**
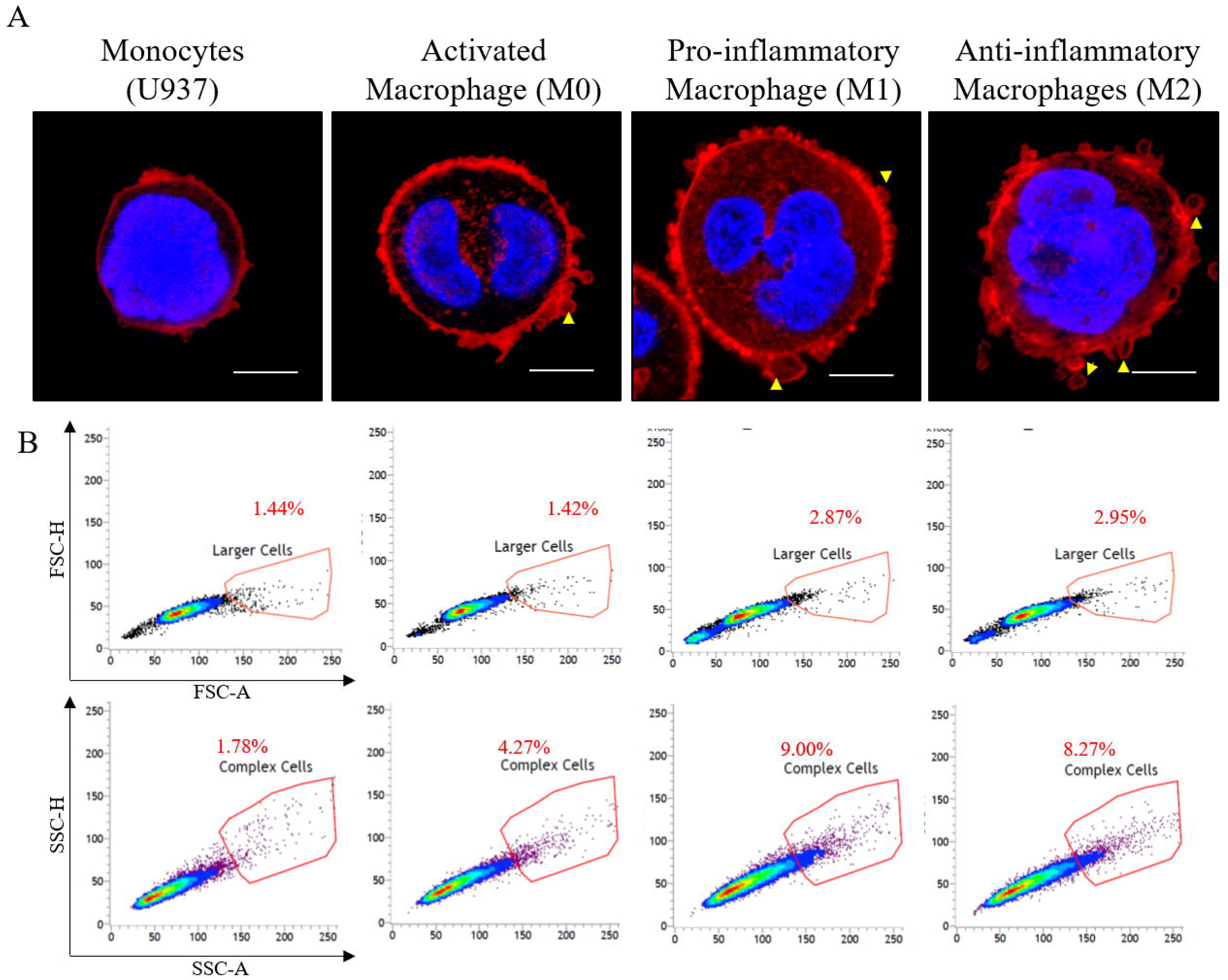
Morphological analysis of monocytes, activated (M0) and polarized (M1 and M2) macrophages generated from monocytic cell line (U937) through CLSM and Flow cytometry. A: Cells were seeded onto glass coverslips and fixed and permeabilized. Nucleus and actin cytoskeleton staining were done with DAPI (blue) and TRITC conjugated phalloidin (red). Multi-nucleation observed in activated and polarized macrophages. Yellow arrow heads denote the macropinocytes made up of actin cytoskeleton. Imaging was done with 63x objective of Nikon AR1 CLSM. Scale bar is 100 µm. B: Cellular size and granularity were analysed through plotting FSC-A/FSC-H and SSC-H/SSC-A values respectively obtained from fixed monocytes, activated and polarized macrophages using BD LSRFortessa flow cytometer. Singlet population were used for plotting. Polarized macrophages (M1 and M2) exhibits increased size and granularity compared to control (monocytes).

Functional aspects of the differentiated macrophages were further analyzed by investigating their CD14/16 expression profile (Fig 2A). Being a CD14+/CD16-monocyte, the U937 population exhibits a higher expression of CD14, which decreases in the M0 state after the PMA-mediated activation. Both kinds of polarized macrophages (M1 and M2) showed a bias towards the CD14^low^-CD16^low^ population, which could be categorized as a subtype of the M2 (M2a) population (Oates et al., 2023). Analysis of pro- and anti-inflammatory cytokines were analyzed through Cytokine Bead Arrays (Fig 2B). Expression of IL-1β, which is a known inflammatory cytokine (Boraschi, 2022) observed to be significantly high among the M1 population, supporting its pro-inflammatory nature. Expression of recognized anti-inflammatory/resolving cytokines such as IL-4 and TGF-β is heightened in the M2 population along with IL-10, which is also reported to have pro-tumorigenic functions (Mannino et al., 2015). Seeing the expression of IL-4, IL-10, and TGF-β together, it can be presumed that the M2 population here exhibits a clear anti-inflammatory profile.

**Figure 2:**
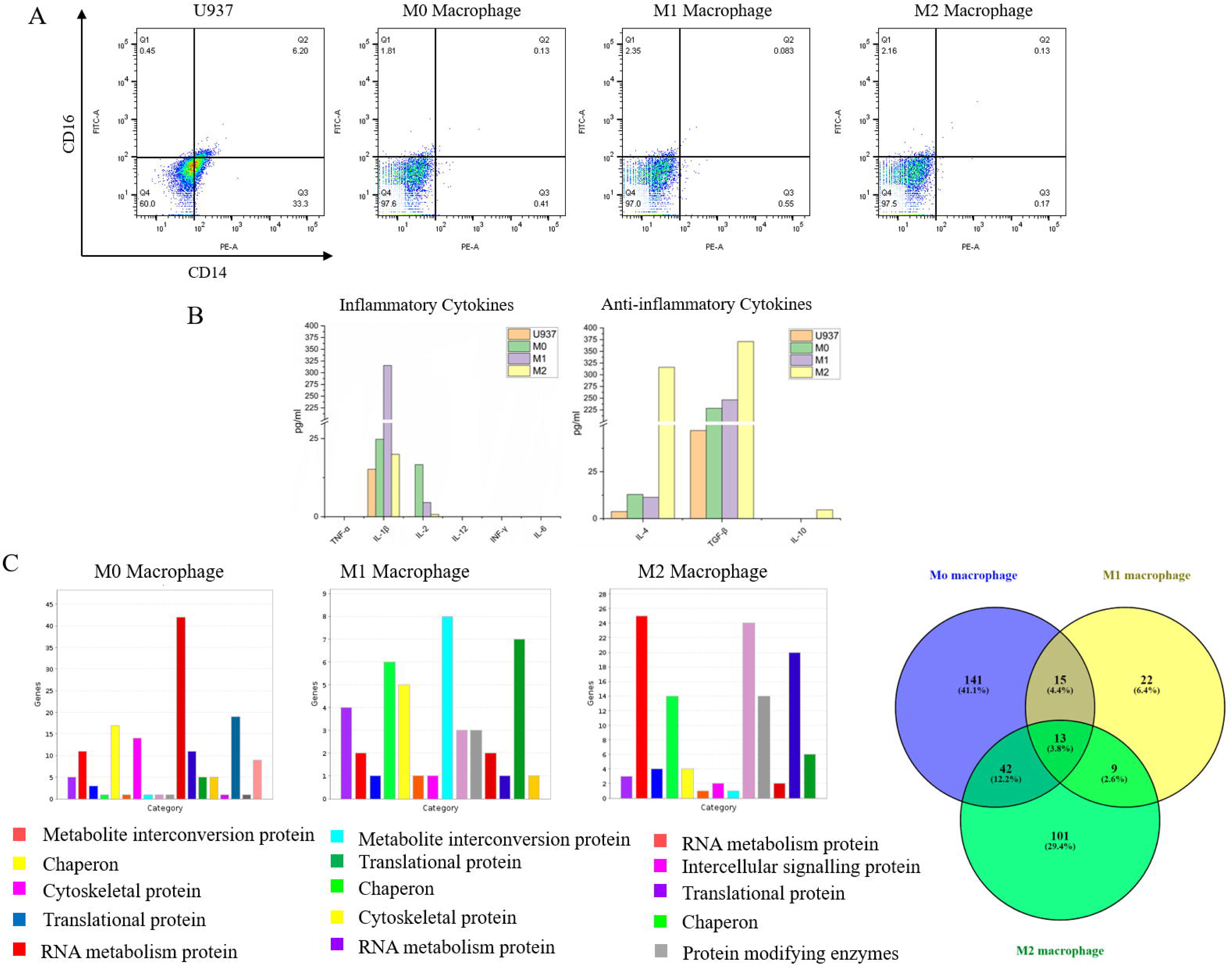
Quantitative analysis of surface markers, inflammatory nature and total proteome of monocytes, activated (M0) and polarized (M1 and M2) macrophages. A: Expression of CD14/CD16 markers on cell population was analysed through Flow cytometry. Cells were fixed and incubated with PE conjugated Anti-Human CD14, FITC conjugated Anti-Human CD16 antibodies. Singlet population was selected for further analysis of CD14/16 expression. B: Expression of pro-and anti-inflammatory cytokines was analyzed through Cytokine bead array using LEGENDplex™ HU Essential Immune Response Panel (13-plex). Conditioned media (serum free) collected from the monocytes, activated (M0) and polarized (M1 and M2) macrophages were subjected to the assay following the manufacturer’s protocol. Values were derived from the fcs files and standard curves generated from BioLegend’s LEGENDplex^TM^ data analysis software. Plotting was done using origin. C: Total proteome analysis was done through differential labelling with reductive dimethylation (ReDiMe) protocol. Linearized and trypsin digested peptides were treated with heavy formaldehyde (CD_2_O) and analyzed through SCIEX TripleTOF^®^ 6600 LC-MS/MS system. Peptide search and quantification was performed using ProteinPilot software and RefSeq protein databased of *Homo sapience*. Monocytes proteome were used for normalization. Upregulated protein candidates (p<0.05) were used for protein class analysis using PANTHER database. Common protein candidates were identified through Venny 2.1 server.

Comparative proteomics analysis of activated and differentiated macrophages further highlights the change in cellular protein level (Fig 2C). We compared the protein signature of activated macrophages (M0) with monocytes and got ∼212 candidates with significant p value (<0.05). When polarized M1 and M2 protein signatures are compared with M0 population, we got ∼60 (M1) and ∼165 (M2) proteins upregulated with significant p value (<0.05) (SI Table 1). When plotted using PANTHER database (Mi and Thomas, 2009), the upregulated proteins from the M0 population belongs to the metabolical pathways, chaperons, cytoskeletal protein, transcriptional and translational family as reported earlier (Sintiprungrat et al., 2010; Minafra et al., 2011). Upregulation of the cytoskeletal protein family also supports the change in cellular shape and morphology. Our study shows the impact of the polarization of M0 into M1 by increasing the expression of metabolical pathways, translational machinery and chaperons among others. M2 population shows upregulation of RNA metabolism, signaling protein, translational machinery and chaperons. Proteins relevant to transporters (He et al., 2018), phagocytosis (Law et al., 2009) are also observed to be upregulated in polarized population. Increased expression proteins involved in transcription, translation, and post-translational modifications highlight the metabolically active state of M1 and M2 compared to M0. When the upregulated proteins were compared between M0, M1, and M2 population, M0 and M2 subtypes showed an increased similarity (12%) compared to M0-M1 (4%) and M1-M2 (2.5%) subset (Fig 2C). This further highlighted the pro-tumorigenic (pro-M2) nature of the activated macrophages (M0) as reported previously (Nascimento et al., 2022). We observed a limited conversion effect of LPS and IL-4-mediated treatment on M0, as experienced by other groups. An earlier study reports the polarization of activated macrophages from U937 origin, producing 10-15% of cells of the M1 subtype and 20-25% of the M2 subtype (Nascimento et al., 2022). For the recruitment study, the adhered population of activated/polarized macrophages was used after harvesting them through scrapping.

A collagen-based (1%) in vitro platform was developed to mimic the chemoattractant-dependent recruitment of monocyte-derived macrophages to breast cancer spheroids. The fabricated spheroids were embedded into the bottom of collagen gel (∼2mm thick), followed by the seeding of monocytes/macrophages on top of it (Fig 3A). As observed in Fig 3B, among the seeded cells, activated (M0) and polarized macrophages (M2) exhibit significant migration towards the spheroids through the collagen. In control cases (without spheroids), monocytes and M0 macrophages exhibit comparable migration, while M2 exhibits minimal migration (M1>M2) (Fig 3B and SI Fig 2). This highly motile nature of monocytes and activated macrophages was also observed elsewhere (Du et al., 2024), which allows them to infiltrate the protein-rich matrix probably through random migration. However, in the presence of spheroid, the seeded cells predominantly exhibit a directed migration towards the bottom. Among all the conditions, M1 subtype displays minimal distance migrated while both the M0 and M2 subtypes migrated the whole distance (∼2 mm) to reach the spheroid at the bottom (SI Fig 2). The persistent migrators or the number of infiltrating macrophages were comparable in both M0 and M2 subtypes. This observation is also supported by recent reports where activated macrophages are capable of reaching the tumor more efficiently (Zhang et al., 2022; Adebowale et al., 2023). In another study, Du et al. (2024) reported the enhanced migrational capacity of tumor-associated monocytes (educated with organoid-conditioned media) compared to both activated (M0) and polarized macrophages (M1 and M2). A competitive migration assay was designed to address the question of whether the monocytes or macrophages are being recruited by some target-specific pathways or not. As observed in Fig 3B, within 18-24 hrs of incubation, the activated macrophages (red) outcompete all three different subtypes (green) and reach the spheroids at the bottom. Here the breast tumor spheroids are observed to explicitly recruit the activated macrophages in the presence of other subtypes. This supports the hypothesis, which suggests the phenomenon of macrophage recruitment as an active process carried out by the tumor and not a random migration process happening to the recruitable cells.

**Figure 3:**
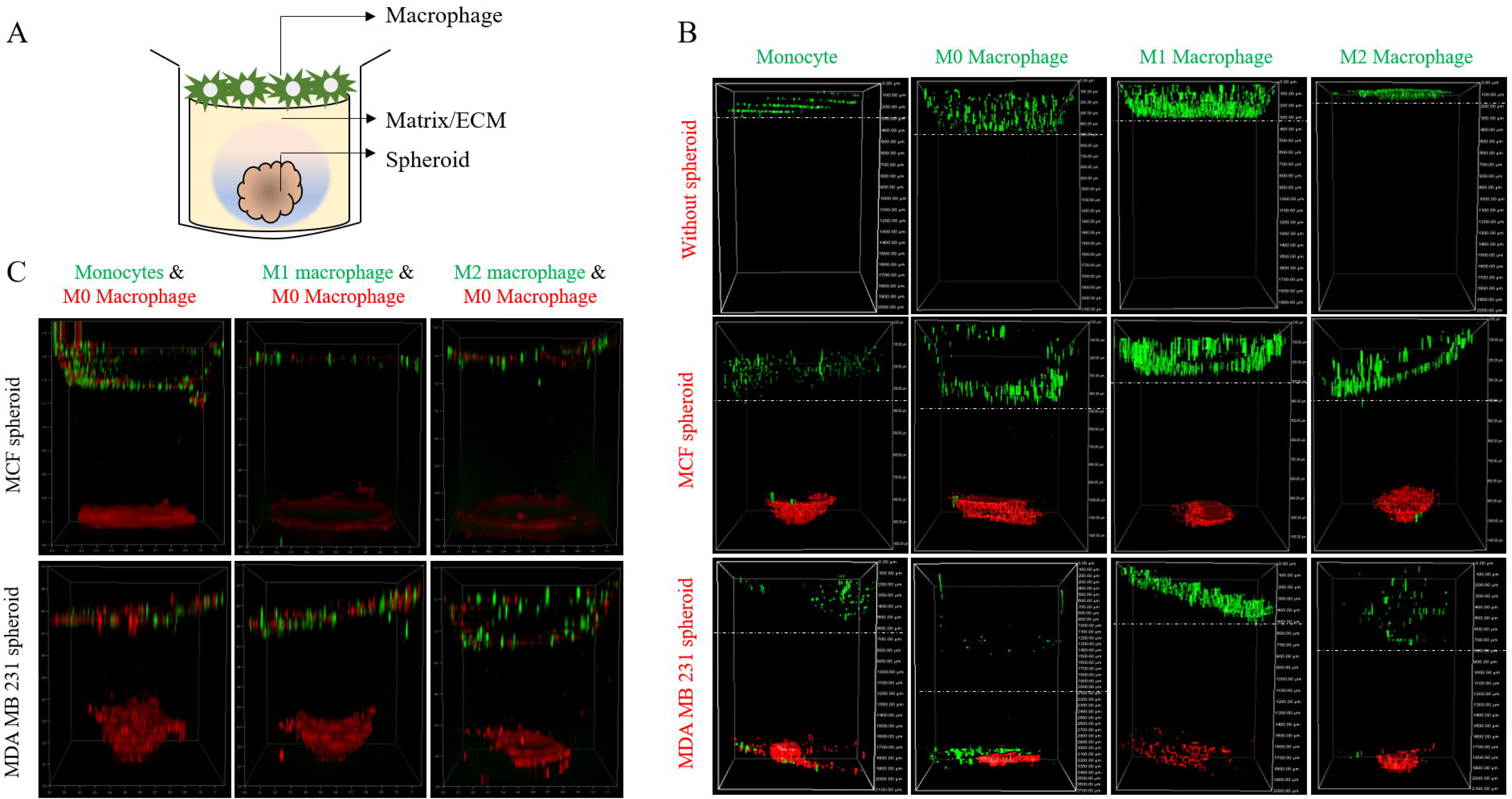
In vitro chemo-attractant mediated macrophage recruitment assay. A: A schematic diagram depicting the entrapment of pre-fabricated pre-stained spheroid into Collagen hydrogel prepared in 96 well plate, followed by the addition of pre-stained monocytes/macrophages on top of it. The thickness of collagen hydrogel would be around 1.8-2 mm. The plate will be incubated in standard cell culture conditions for 16-24 hr followed by CLSM imaging. B: Monoctytes/macrophages pre-stained with CellTracker Green seeded on collagen gel (1%) with and without entrapped spheroid (pre-stained with CellTracker Red) at the bottom. Z-sectioning of collagen layer (1.8-2mm) was done through CLSM after 16-24 hr of seeding. Each optical slice was merged and side profile was generated for the assessment of migration of monocytes/macrophages through the gel. The images were further used to calculate the migratory population (SI fig 2). In control cases, activated macrophages exhibited maximum distance migrated followed by Monocytes and M1 subtypes. M2 macrophages exhibit minimal migration under control set up. However, in presence of spheroids (MCF7 and MDA MB231), all types of immune cells migrated to the bottom of the well, except M1 subtypes, which highlighted the major role of chemotactic migration for macrophage recruitment. C: Competitive recruitment was analysed using two migratory population (Mono vs M0, M0 vs M1 and M0 vs M2) where M0 and Mono/M1/M2 population is pre-stained with CellTracker Red and CellTracker Green respectively. The unstained spheroids of both MCF7 and MDA MB 231 cells were entrapped as before. The imaging and data analysis were done as before. Pre-dominant incorporation of activated macrophages (red) was evident from the spheroid color at the bottom.

As pro-tumorigenic macrophages (M0 and M2) predominantly travel the collagen gel and reach the spheroids at the bottom, we further investigate the ‘physical recruitment’ by retrieving the spheroids and removing the surface-bound macrophages by extensive washing. The flow-cytometric analysis identified the pre-stained (Cytotracker green) monocytes and different subtypes of macrophages incorporated within the spheroids (Fig 4 and SI Fig 3). As observed, M0 (12-15%) and M2 (12%) macrophages exhibit almost similar profiles of incorporation to three types of spheroids, while monocytes and M1 macrophages exhibit minimal recruitment. This preferential recruitment of pro-tumorigenic cells (immune cells of macrophages and monocytic origin) by the tumor has also been reported in recent studies (Zhang et al., 2020; Adebowale et al., 2023; Du et al., 2024). While comparing the incorporation efficiency, we have observed a trend between different types of spheroids (MCF7>MDA MB231>heterotypic), where the most compact and uniform spheroids recruit more cells compared to the aggressive/heterospheroids without any significant difference in their chemokine profile. Among the spheroids, MCF7 ones display high E-cad homodimers, whereas MB MB231 exhibits a lowered expression of E-cad (Manuel et al., 2013). Heterospheroids might exhibit a mixed expression profile of E-cad as the MDA MB231 cells are primarily present on the spheroid surface (SI Fig 1). We presume that the nature of cell-cell adhesion between the cancer cell and the incoming macrophages could be another critical factor to influence the recruitment process as predicted elsewhere (Chae et al., 2018). Though the recruitment process of macrophages into the tumor is not known at either the cellular or molecular level, it would be safe to assume that the spheroids with strong homotypic cell-cell adhesion could be a better target for the physical interaction with macrophages compared to spheroids with hetero-typic interactions. The accessibility of surface receptors might influence macrophage recruitment negatively as it does to breast cancer-associated myeloid-derived suppressor cells (Zhou et al., 2024). It also highlights the importance of macrophage-cancer cell adhesion as the population of finally recruited macrophages probably correlates to the number of adhered macrophages.

**Figure 4:**
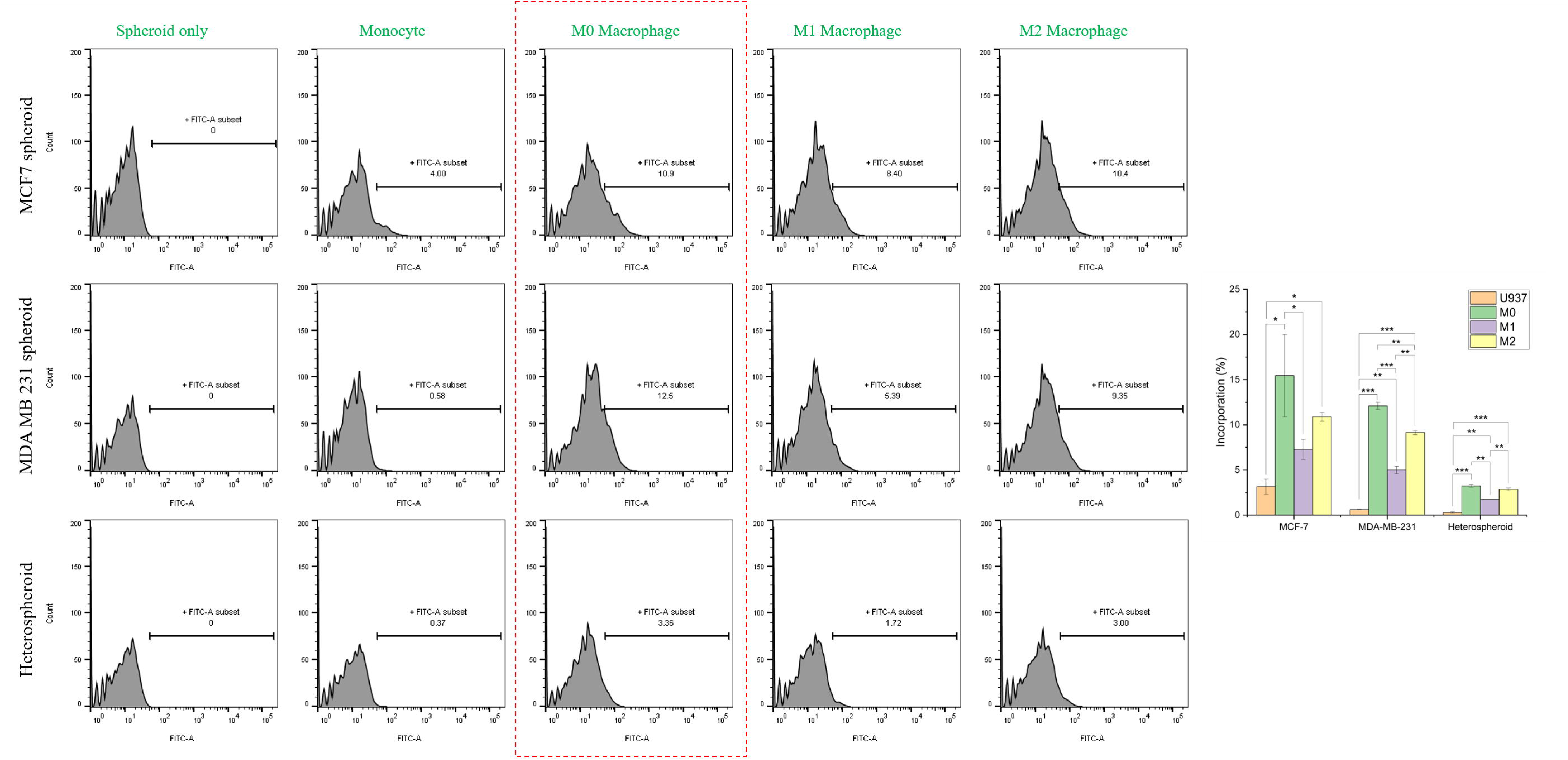
Analysis of recruited monocytes/macrophages through flow-cytometry. The monocyte/macrophage recruitment experiment is repeated with monocytes/macrophages pre-stained with CellTracker Green and unstained spheroids of different types (MCF7, MDA MB231 and heterospheroids) for 16-24 hr. The spheroids were then retrieved from collagen gel and disassociated into single cell suspension. The single cell suspension was subjected to flow cytometric analysis using FSC/SSC cluster, and FITC-A channel gating. The FITC positive cells (monocytes/macrophages) were quantified after singlet analysis (SI Fig 3). Unstained spheroids were used as control. In all three types of spheroids, the incorporation profile is as observed to be as follows, M0>M2>M1>Monocytes. The incorporated cells were plotted using Originpro software. For statistical analysis, One-Way ANOVA was done followed by Tukey’s test (post hoc analysis) for comparison between groups. *P* ≤ 0.05 was considered as statistically significant for all experiments, and values were assigned accordingly (\**P* ≤ 0.05, \*\**P* ≤ 0.005, \*\*\**P* ≤ 0.001).

To understand the target-specific recruitment process, we further assess the chemokine profile of three different kinds of spheroids fabricated with non-metastatic (MCF7), metastatic (MDA MB231), and mixed (MCF7+MDA MB231) breast cancer cells through Cytokine Bead Array against the panel of 13 chemokines. As seen in Fig 5A, all three kinds of spheroids display significant amount of CXCL8, CCL2, CXCL1, CCL20, CCL4, and CCL5. Among the three types of spheroids, MDA MB231 spheroids showed increased production of CXCL8, CXCL10, CCL1, and CCL20 compared to others. The expression profile is comparable with breast cancer chemokine profiles as reported in earlier studies (Svensson et al., 2015; Liu et al., 2020; Masih et al., 2022), along with other solid tumors (Baier et al., 2005), with increased expression of CCL2, CCL4, CCL20, and CXCL8. Among them, the CCL2/CCR2 axis has been widely studied for the recruitment of tumor-associated macrophages. Multiple reports have identified this pathway as a potential therapeutic target in order to minimize TAM recruitment (Qian et al., 2011; Chen et al., 2017). However, the clinical failure of CCL2 targeting therapeutics (CNTO888, MLN1202) highlighted the complexity of tumor-mediated recruitment of TAMs, along with the possibility of other pathways. Along with CCL2, other chemokines, such as CXCL8 (Xiong et al., 2022), CCL20 (Boyle et al., 2015), and CCL5 (Walens et al., 2019), are also highlighted for their role in TAM recruitment. They are identified as potential therapeutic targets (Kitamura and Pollard, 2015; Argyle and Kitamura, 2018). As the CCL2/CCR2 axis of TAM recruitment has been studied over without much success, we further checked the probability of other chemokine mediated pathways playing critical roles in TAM recruitment by investigating their receptor status within the macrophage population (Fig 5B). As observed, mRNA levels of CCR6, CCR3, and CXCR1 exhibit significant increments in the M0 population except for CCR5, which is high in the M2 population. To correlate the in vitro data and TAM recruitment in tumors, we further crosschecked the relation between the macrophage infiltration and expression of the following chemokine receptors using the TIMER 2.0 server (Li et al., 2020). The purity-adjusted Spearman’s rho was calculated using the TIMER algorithm, which indicates a positive correlation between the expression level of CCR6, CCR3 and CXCR1 and tumor infiltration rate of macrophage (Fig 5C). These findings are in line with other reports where CXCL8/CXCR1/2, CCR20/CCR6, CXCL1/CXCLR1 and CCL3/CCR3 axis, are identified as auxiliary pathways (Boyle et al., 2015; Nandi et al., 2016; Xiong et al., 2022) and could act as the potential target for therapeutic intervention.

**Figure 5:**
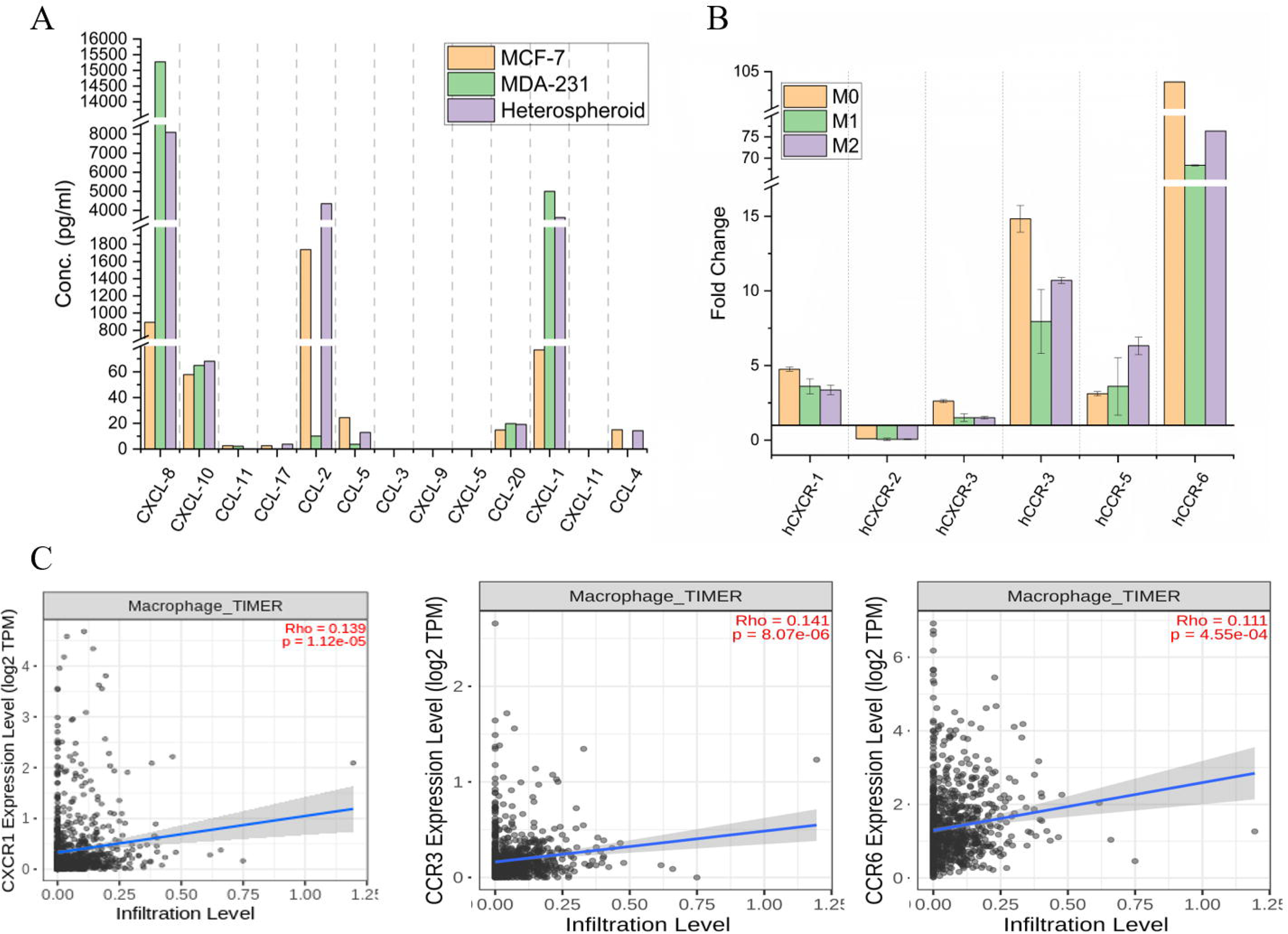
Analysis of chemotactic interaction between the spheroids and monocytes/macrophages and their role in macrophage recruitment. A: Conditioned media collected from three types of spheroids (MCF7, MDA MB231 and heterospheroids) were analysed using LEGENDplex™ Hu Pro-inflam. chemokine panel (13 plex) cytokine bead array. All three spheroids types express CXCL8, CXCL11, CCL2, CCL5, CCL20, CXCL1 and CCL4. No difference between the spheroid expression level except for CXCL8 and CCL2 are observed. B: Expression level of chemokine receptors on M0, M1 and M2 population was analysis through QPCR. The fold change values (Mean±SE) (normalized with monocytes) were plotted using Origin pro software. mRNA profile of hCXCR1, hCXCR3, hCCR3, and hCCR6 are observed to be high in M0 compared to other subsets of macrophages except hCCR5. C: Correlation analysis of CXCR1, CCR3 and CCR6 expression and macrophage infiltration through TIMER2.0 show positive correlation with macrophage infiltration in breast tumor.

After the recruitment, the macrophages presumably go through an overhaul of their transcriptomic and transcriptional profile and finally can be categorized as Tumor-associated Macrophages (Cassetta et al., 2019; Du et al., 2024), which should happen through physical contact with tumor cells or tumor-associated stromal cells (Gok et al., 2019).

Here, we want to investigate the impact of the recruited macrophages on spheroid viability and migration. We used MCF7 spheroids to assess the cytotoxic effect of macrophages as they incorporated most of the macrophages. The control and retrieved spheroids with incorporated M0 macrophages were allowed to incubate for 72 hrs under standard conditions on agarose-coated plates. Fig 6A displays no deleterious effect observed in the context of the dead cell population (Calcein AM/PI staining). Here, incorporated macrophages (15% of the total population) do not exhibit any cytotoxicity towards the spheroids.

**Figure 6:**
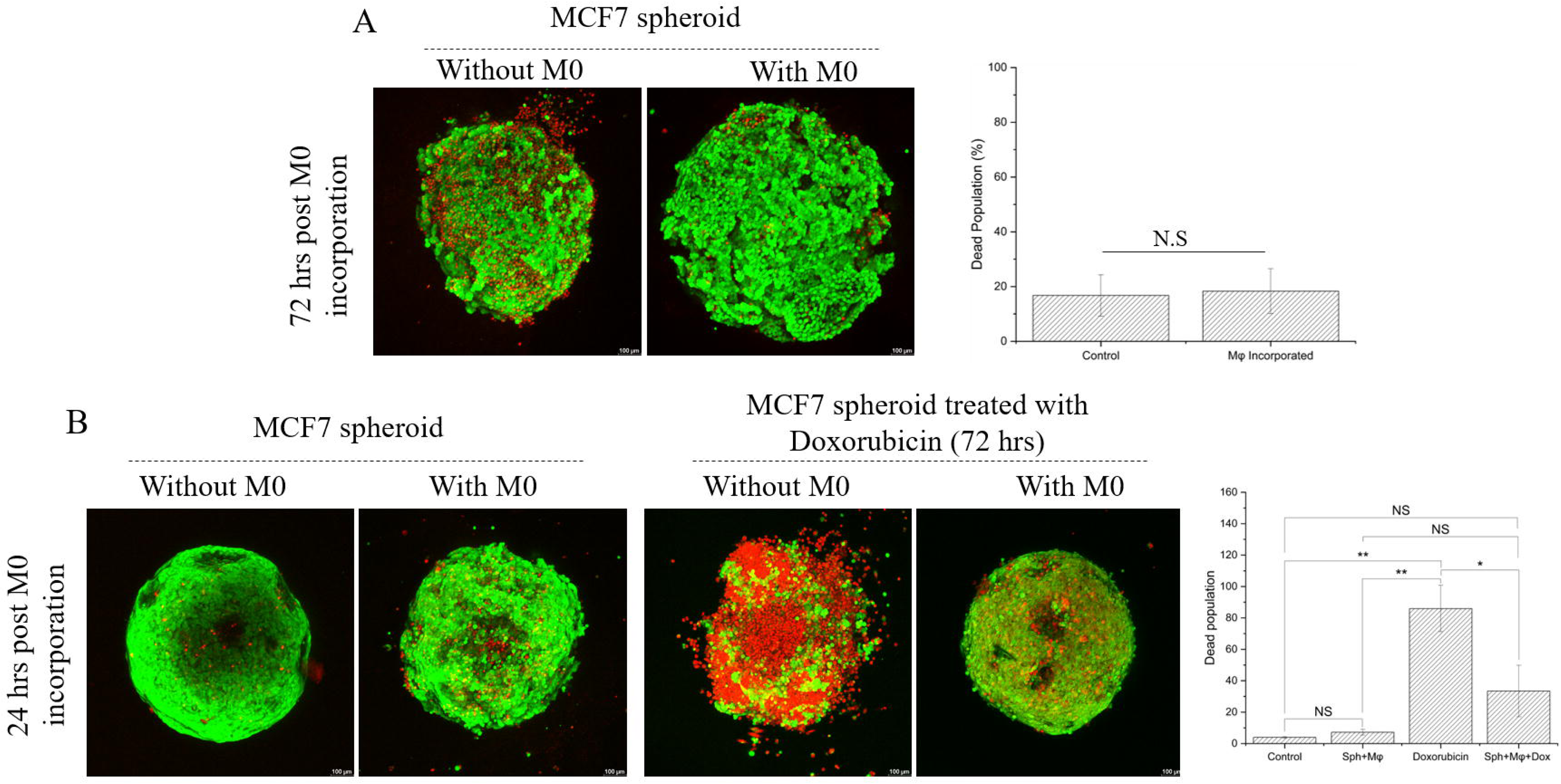
Analysis of recruited macrophage mediated cytotoxicity and drug resistance on spheroids. A. Control and M0 incorporated spheroids were incubated under standard cell culture conditions for 72 hr and analyzed for cell death using Calcein AM and PI staining. Samples were analyzed by CLSM. Mean fluorescent intensity of PI was measured and plotted. No significant difference is observed between the samples. Scale bar is 100µm. B: Control and M0 incorporated spheroids were subjected to Doxorubicin (50µM) treatment for 72 hrs followed by staining with Calcein AM/PI staining. Mean fluorescent intensity of PI was measured and plotted. Spheroids with M0 exhibited significant resistance to the drug.

Further, we want to test the role of the incorporated macrophages in the development of drug resistance among the above-mentioned spheroids. As observed (Fig 6B), spheroids with and without Macrophages subjected to chemotherapeutic treatment (50 µm of Doxorubicin) for 72 hrs exhibit a clear distinction in alive/dead cells. Control and treated spheroids stained with Calcein AM/PI, showed significantly high PI staining in no-M0 spheroids compared to the spheroids with M0 macrophages. Though it’s not clear how TAMs (U937 derived) induce drug resistance, they probably follow the common IL6-mediated pathway (Dong et al., 2020a) or through exosomes (Dong et al., 2020b), which needs to be investigated further.

Additionally, we want to check the influence of the incorporated macrophages on the migration efficiency of the spheroids, as TAMs reportedly enhance the metastatic potential of solid tumors (Larionova et al., 2020). To assess the migration potential, we retrieved the control and M0 (U937 derived) incorporated spheroids with different metastatic ability (MCF7, MDA MB231 and Heterospheroid) and embedded them in Collagen type 1 (1%) gel for 24-72 hrs. As demonstrated in Fig 7, the control MCF7 spheroids (mCherry tagged) inflated over time (72 hrs) without any visible migratory cells. After the incorporation of M0 macrophages, MCF7 spheroids do not exhibit much improvement in migration patterns except for a few individual cells in the nearby ECM. However, the MDA MB231 spheroids (eGFP tagged) with M0 macrophages exhibit more sprouting cells even at 24 hrs compared to its counterpart. Over time (72hrs), the number of spindle-shaped migratory cells increased rapidly in both cases. However, the initially migrated population increased the total migration efficiency in M0 macrophage-incorporated spheroids within the nearby ECM. While the monotypic spheroids remain true to their inherent migration capacity, the heterospheroids exhibit few migratory MDA MB231 cells in control samples. The migration pattern of heterospheroids differs drastically in the M0 macrophage incorporated condition with time. Though some MDA MB231 cells are observed in the local vicinity, the majority of them are observed to travel far compared to other conditions with few clustered MCF7 cells (SI Fig 4).

**Figure 7:**
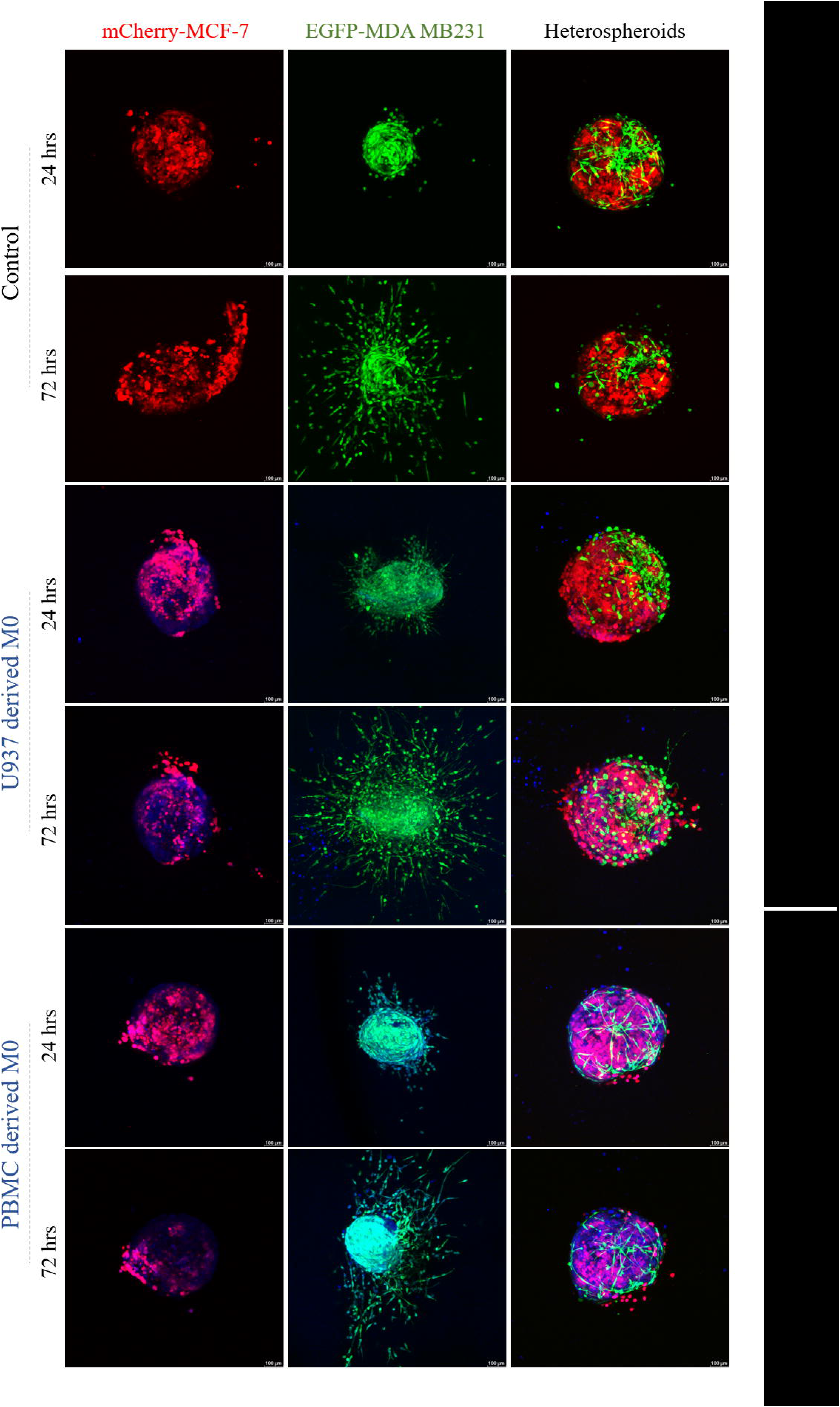
Impact of incorporated macrophages of U937 and human monocyte origin on proteolytic migration of different types of spheroids. Spheroids of monotypic (MCF7-mCherry, MDA MB231-EGFP) and heterotypic origin were used for recruitment of M0 macrophage (of U937 and human monocyte origin) pre-stained with Celltracker Blue as before. M0-recruited spheroids were retrieved and entrapped in Collagen (1%) hydrogel before further analysis. Imaging of same spheroids was done through CLSM (10x objective) for 24 hr and 72 hr through Z-sectioning. Maximum projection was done for post-image analysis. Scale bar is 100µm. Extensive migration of MDA MB231 cells (green) from heterospheroids was analyzed through Tile mode imaging (SI Fig 4).

Furthermore, the impact of M0 macrophages originating from healthy monocytes (human) on spheroid migration in the collagen was tested. Human monocyte-derived M0 macrophages repeat the localization and migrational pattern as U937-derived macrophages. It should be noted that the incorporated macrophages can only enhance the metastatic behavior of the spheroids without drastically altering them. Interestingly the localization of incorporated macrophages also differs between different types of spheroids. In MCF7 spheroids, the macrophages (blue) are observed on the surface of the spheroids, while in MDAMB231 and heterotypic ones, they appeared to be entered into the core (Fig 7).

We assume that the localization of the TAMs and their physical interaction with cancer cells play a significant role in the activation of their pro-tumorigenic activities. However, other reports also suggest that the occurrence of TAM-like capabilities can happen through soluble factors (Chen et al., 2018). To confirm the role of physical interaction on TAM generation, we have also generated tumor-educated macrophages (TEM) by exposing the M0 population to the conditioned media collected from the spheroids (MCF7 or MDA MB231). The conditioned media collected from those TEMs were used to analyze the migration pattern of MCF7 and MDA MB231 spheroids within collagen gel (1%) (SI Fig 5). No significant difference between the control and treated spheroids in terms of migration efficiency was observed in our study, which supported the importance of physical interaction between the cancer cells and recruited macrophages.

Additionally, we want to confirm the nature of the incorporated macrophages by assessing the expression of one of the recently reported breast tumor TAM markers, namely Siglec-1 (CD169), as the expression of conventional TAM marker CD163 becomes contradictory (Cassetta et al., 2019). Siglec-1 is a surface marker that interacts with Sialic acid residues on target cells and influences macrophage-mediated phagocytosis. In earlier reports, Siglec-1 expression was observed to be limited to TAMs and not in circulatory monocytes (Cassetta et al., 2019), which is reflected in our study too (Fig 8A), where the mRNA profile of Siglec-1 is highest in activated and polarized macrophages but not in monocytes. Expression of Siglec-1 seems to lowered in polarized and conditioned media-educated macrophages (TEM1 and TEM2), which highlighted the difference between the polarized and tumor associated macrophages. To support the point of considering Siglec-1 as a TAM marker, we further crosschecked its present within the M0-incorporated spheroids. Presence of Siglec-1 mRNA is observed to be upregulated in M0-incorporated spheroids. Additionally, IHC of Siglec-1 done with the breast tumor (Indian patient) sample shows positive for Siglect-1+ (Fig 8C). Presence of Siglec-1positive macrophages are reportedly connected to both anti-tumor (Shiota et al., 2016) and pro-tumorigenic activities (Jing et al., 2020) in breast tumor. Increased expression of Siglec-1 protein is found to be positively correlated with the infiltration efficiency of macrophages within different subtypes of breast tumors (including basal, Her2+, Luminal A and B) when analyzed using TIMER 2.0 web server (Fig 8B and SI Fig 6). This highlights the positive role of Siglec-1 with macrophage recruitment into the tumor, probably through an Integrin-mediated path (SI Fig 7) (Szklarczyk et al., 2023). As the expression of Siglec-1 is found to be highly relevant to tumor-mediated macrophage recruitment, their role in macrophage recruitment and activity should be assessed further.

**Figure 8:**
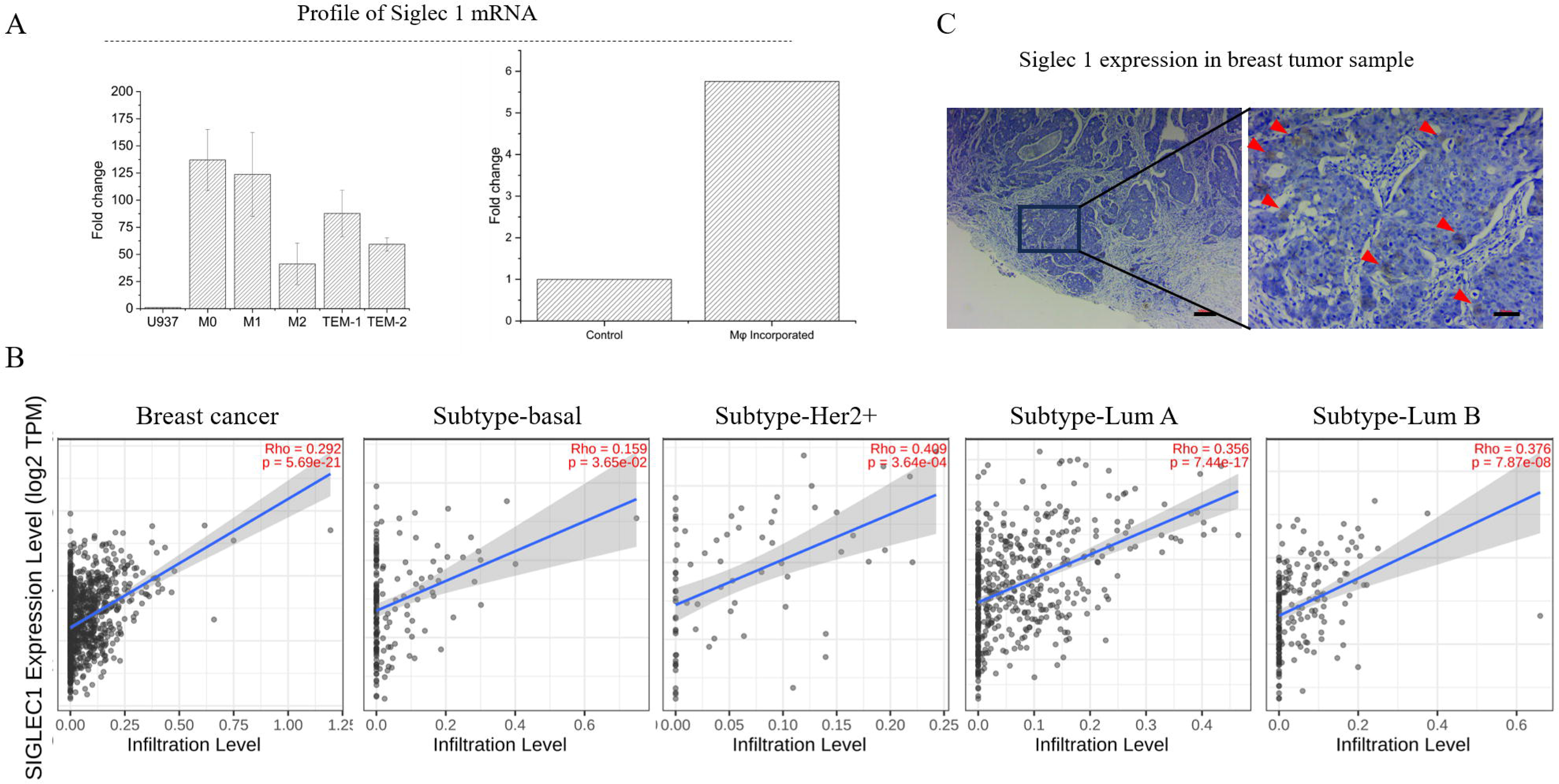
Analysis of TAM marker Siglec-1 expression and its correlation with macrophage infiltration. A: mRNA profiling of Siglec-1 (CD169) was done on U937, M0, M1, M2 and tumor educated macrophage subtypes (TEM1 and TEM2). Activated macrophages showed highest mRNA level compared to other subtypes. Siglec-1 mRNA is also abundant in spheroids containing M0 macrophages compared to control spheroids. B: Siglec-1 expression is positively correlated with Macrophage infiltration among different types of breast cancer (basal, HER2+, Luminal A and B). Expression of Siglec-1 is also correlated with macrophage infiltration analyzed through EPIC, TIMER and XCELL (SI Fig 6). C: Siglec-1 expression is also observed in breast tumor (Indian patient) through Immunohistochemistry. The red arrows denoted the Siglec-1 expression.

Conversion of incorporated macrophages into TAMs probably required the participation of multiple cytokines and growth factors secreted by the tumors. To identify the probable candidates, we screen the cytokines and growth factors secreted by the MCF7 and MDA MB231 spheroids (SI Fig 8). Expression of IL-6, VEGF, PDGF-AA and M-CSF in the secretome is observed. Diverse role of IL-6 in macrophage recruitment is highlighted in recent work (Du et al., 2024). M-CSF originated from tumor also reported to influence TAM behaviour (Yi et al., 2024). However, it can be assumed that the physical interaction between the tumor and recruited macrophages also play a critical role in TAM conversion along with the secretory factors as CM treated macrophages didn’t exhibit complete conversion. Further probing into the Tumor cell-Macrophage interaction is required to understand the phenomenon.

In recent years, few reports suggested the critical role of pro-tumorigenic macrophages (M2) (Dwyer et al., 2016; Hacker et al., 2023; Mazan and Marusiak, 2024) and monocytes (Du et al., 2024) on tumor-mediated recruitment process, we believe that our report is the first one to describe the pro-tumorigenic role of activated macrophages and their influence on spheroid behavior.

## Conclusion

The interaction between tumors and immune cells is critical as they influence tumor biology and behavior in the context of tumor survival and invasion. Early-stage tumors tend to develop an immune-suppressive environment using the pro-tumorigenic components of immune cells such as macrophages. Developing a cost-effective in vitro platform to assess the crosstalk is vital for many reasons. We find that both mono and heterotypic breast tumor spheroids preferentially recruit activated macrophages through chemokine-dependent pathways. Furthermore, the incorporated M0 macrophages exhibit pro-tumorigenic capabilities and express potential TAM markers. The in vitro platform can be further used to assess the cancer-immune cell interaction at close proximity. The spheroid-based platform has the potential to be a high-throughput assay for screening cell-based immunotherapeutics in the future.

## Materials and Methods

### Materials

Basic cell culture media like DMEM, RPMI-1640, Fetal Bovine Serum, TPVG, Penicillin/ streptomycin, HiSep™, Triton X-100, BSA and Doxorubicin-HCL were procured from HiMedia, India. Phorbol (PMA) and LPS was procured from Sigma Life Science. Growth factors and cytokines such as M-CSF and IL-4 were obtained from Sino Biologicals, CellTracker live cell stains (CellTracker Red, CellTracker Green and CellTracker Blue), Rhodamine conjugated Phalloidin, DAPI, Calcein-AM, Propidium Iodide, Rat tail Collagen type I, Trizol, Revertaid First Strand cDNA synthesis kit, and PowerUp SYBR Green Master Mix, Lipofectamine3000 were procured from Thermo scientific. Transfection agent, Virafect was acquired from Promega. Primers used for semi quantitative and quantitative PCR were procured from Eurofin, India.

Antibodies (Human Siglec-1, PE conjugated Anti-Human CD14, FITC conjugated Anti-Human CD16) were obtained from Thermo Fisher Inc. Accutase was purchased from Gibco. LEGENDplexTM Hu Growth factor Panel (13-plex), LEGENDplex™ HU Essential Immune Response Panel (13-plex) and LEGENDplex™ Hu Pro-inflam. chemokine panel (13plex) were procured from Biolegend for Cytokine bead array (SI Table 2, SI Table 3, SI Table 4). pmCherry C1 vector was procured from Takara. pLenti-cmv-gfp-puro cloning vector (a generous gift from Prof. Eric Campeau & Prof. Paul Kaufman) packaging plasmid (psPAX3) (a generous gift from Prof. Didier Trono) and envelope expressing plasmids (pMD2.G) (a generous gift from Prof. Didier Trono) were received from Addgene. All the Cell culture grade Plasticware was procured from Tarsons, India.

#### Procurement of cell lines and their maintenance

Cell lines such as MCF7, MDA-MB-231, HEK 293T, and U937 cells were procured from NCCS Cell Repository, Pune, India. MCF7, MDA-MB-231 and HEK 293T cells were maintained in high glucose DMEM and U937 cells were maintained in RPMI-1640 supplemented with 10% FBS and Penicillin-streptomycin under standard cell culture conditions. MCF7, MDA-MB-231, HEK 293T and U937 cells were harvested after reaching 70-80% confluency. Adherent cells like MCF7, MDA 231 and HEK 293T were collected after trypsinization, while non-adherent U937 cells were harvested by centrifugation. All the centrifugations were carried out at 1200rpm for 5mins at room temperature.

Blood from healthy human volunteers were collected from the local blood bank following the approval of Institute Ethics Committee for human PBMCs. Isolation of PBMCs were done following the protocol stated elsewhere. Briefly, the buffy coat was diluted with equal volume of HBSS and overlaid on top of 20ml of HiSep™. Post-centrifugation the WBC layer was collected in HBSS and washed twice. The cell pellet was resuspended in RPMI basal medium and seeded in 100mm tissue culture plate. After 24 hrs. non-adhered cells were discarded and the adhered monocytes were washed twice with HBSS. Monocytes were harvested by gentle scraping and used for further experiments.

#### Development of mCherry-MCF-7 and EGFP-MDA-MB-231 cells

MCF-7 cells were transfected with pmCherry-C1 vector using ViraFect^TM^ following manufacturer instructions. Briefly, 1μg pmCherry-C1 plasmid DNA was mixed with 3μl of transfection reagent (1:3 DNA: reagent ratio). The mixture was kept at room temperature for 20mins and was added to MCF-7 cells (50,000/well) followed by 24-48hrs incubation. Transfected cells were selected using G418 (800μg/ml) containing complete DMEM medium. A single colony was picked by localized trypsinization and maintained in G418 containing media for weeks.

pLenti-CMV-GFP-Puro vector was used to develop EGFP-MDA MB231 cell line in Biosafety level 2 culture facility following the Institutional Biosafety Committee approval. HEK-293T cells were transfected with PAX2 (1.3pmol), pMD2.G (0.72pmol) and pLenti-CMV-GFP-Puro (1.64pmol) using Lipofectamine 3000 reagent (1:3 DNA:Plasmid ratio) following the manufacturer’s instructions. The culture supernatant containing the viral particles was collected and filtered through 0.45μ filter. Confluent culture of MDA-MB-231 was infected with 0.5ml viral supernatant supplemented with 8μg/ml polybrene. Cells were thoroughly washed with PBS after 24 hrs. and checked for GFP expression under fluorescence microscope. Transduced cells were selected with 1μg/ml Puromycin for 1week.

Both kinds of stably transformed cells (mcherry-MCF7 and EGFP-MDA MB231) were sorted using BD FACS Aria cell sorter. Sorted population of MCF-7 pmCherry^Hi^ and MDA-MB-231 eGFP^Hi^ cells were further cultured and used for experimentation.

#### Differentiation and polarization of U937 cells into activated (M0) and polarized (M1 and M2) macrophages

U937 cells were differentiated to activated macrophages by following the method described earlier (Kuno et al., 2020; Lund et al., 2016). Briefly for differentiation, 1X10^5^ cells/ml monocytes (U937 cells) were stimulated with 100nM PMA for 24hrs followed by recovery in complete medium for 72hrs. Such activated macrophages (M0) were further treated with either 100ng/ml LPS or 20ng/ml IL-4 for 48hrs to polarize them into M1 and M2 macrophages respectively. The macrophage morphology was observed by phase contrast microscopy (Magnus).

#### Characterization of monocyte (U937) derived macrophages (MDMs)

##### Analysis of size and granularity

Extent of morphological changes during differentiation and polarization of U937 derived macrophages (M0, M1 and M2) was measured by assessing light scattering properties in flow cytometer (BD FACSVerse). Singlets analysis was done by plotting FSC-H vs FSC-A values. Cellular complexity was further analyzed by plotting SSC-H vs SSC-A values.

##### Sub-cellular structures

Monocyte derived naïve (M0) and terminally differentially macrophages (M1 and M2) were seeded and differentiated on coverslips. Upon differentiation cells were fixed with 4% PFA for 10mins at room temperature. The cells were washed thrice with PBS following permeabilization with 0.1% Triton X-100. The coverslips were blocked in 5%BSA and stained with Rhodamine conjugated phalloidin (Thermo)(1:100 dilution) for 1hr at room temperature. Nuclei were counterstained with DAPI (Thermo) (1:500 dilution) for 10mins at room temperature. The samples were imaged using CLSM (Nikon AR1) under 63X objective.

##### Profiling of CD14 and CD16 expression after activation and differentiation

U937 cells were harvested by centrifugation and U937 derived macrophages (M0, M1 and M2) were harvested by gentle scrapping. Cells were fixed in 2%PFA for 20mins at room temperature followed by the wash and blocking in 3% BSA for 1hr at room temperature. The cells were then incubated with Anti-CD14 antibody conjugated with PE (20μl reagent per 10^6^ cells) and Anti-CD16 antibody conjugated with FITC (20μl reagent per 10^6^ cells). Cells were washed thrice with PBS and resuspended in 200μl of PBS and the data was acquired in flow cytometer (BD LSRFortessa). Analysis was restricted to singlet cells identified by SSC-A vs FSC-A plot. FSC-H vs FSC-A was used to identify singlets. Expression of CD14 and CD16 was assessed by checking PE and FITC signals respectively.

##### Analysis of total proteome of MDMs

Monocytes and macrophages (M0, M1 and M2) were harvested by gentle scrapping and lysed with cell lysis buffer containing 9M urea made in 50mM triethylammonium bicarbonate (TEAB) buffer and were subjected to probe sonication. Further, cell lysates were centrifuged at 1000 rcf for 5mins for removal of cell debris. Supernatant was collected in fresh tube & was subjected to reduction of disulfide bond by treating with 300mM K_2_CO_3_ & 100mM tris-2-carboxyethy-phosphine (TCEP) followed by alkylation with 20mM Iodoacetamide (IAA) at least for 16hrs. The linearized proteins were digested in solution with sequencing-grade modified trypsin in 1:50 ratio. Post trypsin digestion, peptides were subjected to differential labeling with reductive dimethylation (ReDiMe). During ReDiMe, the peptides obtained from monocytes were treated with light formaldehyde (CH_2_O) giving a light label and the peptides obtained from each of the subtypes (M0, M1 and M2) were treated with heavy formaldehyde (CD_2_O) giving a heavy label. Post ReDiMe labelling, the heavy and light labelled peptides were mixed and desalted using an established stage-tip protocol containing C18 resin (Rappsilber et al., 2007). The desalted samples were then subjected to mass spectrometry in SCIEX TripleTOF^®^ 6600 LC-MS/MS system for comparative proteomic analysis using previously reported run conditions. Briefly, the proteomics samples were acquired using an information-dependent acquisition (IDA) method over an m/z range of 200 - 2000. A full MS survey scan was conducted followed by MS/MS fragmentation of 15 abundant peptides detected in the MS survey scan. Additionally, in order to increase the peptide coverage a dynamic exclusion was also enabled (repeat count = 2; exclusion duration = 6 seconds). The proteomics analysis of three biological replicates was used for the further analysis. The peptide search and quantification were performed using ProteinPilot software (version 2.0.1, SCIEX). The search database used for the study was RefSeq protein database of *Homo sapiens* (Release 108, last modified on 5th May 2016). In all the searches, iodoacetamide alkylation of cysteine was described as static modification and oxidation of methionine and N-terminal acetylation were described as variable modifications. Additionally, ReDiMe algorithm was selected for quantification of identified peptides. In all the peptide searches, the mass tolerance of MS and MS/MS ions was placed at 20 and 50 ppm respectively. Furthermore, the peptides were also searched against a decoy database by the software to calculate false discovery rate (FDR) which was specified to <1% to filter falsely identified peptide and proteins. Significantly upregulated proteins from technical replicates (with p < 0.05) were selected for further analysis. The role of differentially expressed proteins in different cellular process, pathway and cellular compartmentalization were analyzed using PANTHER Pathway server (Mi and Thomas, 2009). Upregulated proteins were further plotted as per their protein class and top five classes are mentioned in the figure. Commonality of significantly upregulated proteins from M0, M1 and M2 population were analyzed using Venn diagram using Venny 2.1 server (Oliveros. 2007-2015).

##### Expression profiling of pro-inflammatory cytokines

Differentiated macrophages were washed with PBS and incubated with 3ml of serum free medium for overnight. The conditioned medium was collected for secretome analysis using the Human Essential Immune Response Panel (13-plex) (Biolegend) designed for Cytokine Bead Assay following manufacturer’s protocol. Briefly, 25μl of conditioned medium was diluted with equal volume of assay buffer. 25μl of capture beads were added to the dilute conditioned medium and incubated at room temperature with gentle shaking for 2hrs. Beads were separated by centrifugation and washed once with washing buffer followed by resuspension and incubation with 25μl of biotinylated detection antibodies for 1hr at room temperature with shaking. Finally, 25μl of detection reagent (streptavidin conjugated with PE) is added to the mixture and incubated for 30mins at room temperature with shaking. Following separation of beads and a wash, the beads were resuspended in 150μl of wash buffer and readings were obtained on a flow cytometer with appropriate assay settings. The fcs files were analyzed using BioLegend’s LEGENDplex^TM^ data analysis software. The MFI values of samples were used to quantify the analytes using standard graph of respective analyte.

#### Fabrication of mono-and hetero-spheroids of breast cancer cells

Individual spheroids of MCF7 and MDA MB231 cells were fabricated using agarose overlay method described elsewhere (Gayan et al., 2017). Briefly 70-80% confluent cells were harvested and 6000 cells/well were seeded on 1% Agarose (Lonza) coated 96well plates. The plates were then centrifuged at 2000rpm for 5minutes at room temperature using swing bucket rotor. These spheroids were allowed to settle for 2days, post which Collagen-I was added to the final concentration of 6μg/ml per well in 100μl of medium and the plate was centrifuged at 500g for 15mins. The plates were kept in the incubator at 37°C for 5-7days. In case of Heterospheroids, 3000 cells of MCF7 and 3000 cells of MDA MB231 were mixed together and were seeded on these agarose coated 96well plates for two days, post which collagen-I was added to the final concentration of 6μg/ml per well in 100μl of medium and the plate was centrifuged at 500g for 15mins followed by incubation of 5-7 days.

Fabrication of monotypic and heterotypic spheroids from fluorescently tagged cells (mCherry-MCF7 and EGFP-MDAMB231) were done as before.

#### Recruitment of Monocytes and monocytes derived macrophages to the spheroids

Monotypic and heterotypic spheroids were fabricated and stained with CellTracker^TM^ Red dye for 45 min-1hr at 37°C. Stained spheroids were deposited on the agarose coated 96 well plates and overlaid with collagen solution (1mg/ml). Polymerization of collagen was mediated by NaOH at 37°C for 30mins. Monocytes and activated macrophages (M0, M1 and M2) were scrapped gently and incubated with CellTracker™ Green dye for 45 min-1hr at 37°C. Followed by the polymerization of collagen, 5000 pre-stained Monocytes/macrophages were seeded on top of the collagen gel and incubated under standard cell culture conditions for 16-18hrs. Imaging was done using Confocal Laser Scanning Microscopy (CLSM) (Nikon AR1) under 10X objective using Z scanning mode. Post-acquisition image analysis was carried out using Nikon NIS software. Automated measurement tool was used to calculate the number of monocytes/macrophages in each optical section. Layer of cells at top of the collagen defined the starting point (0µm). Percent cell population was plotted against distance migrated to analyze migration potential of subset of macrophages. For each experiment 500 randomly selected cells (RAND function, Microsoft excel) were used to calculate mean migrated distance and 100 randomly selected cells were used to represent migration distance of each cell type per experiment.

#### Competitive recruitment of MDMs into spheroids

Similar to previous experiments, here unstained cancer spheroids were deposited on agarose coated 96 well plates and overlaid with 1mg/ml collagen. Upon polymerization 2 different types of prestained macrophages (M1 vs M0, or M2 vs M0 and U937 vs M0) were seeded on top of the collagen and their migration was observed as before through CLSM (Leica TC SP8). Migration of different macrophages was compared with respect to each other.

#### Quantitative analysis of the incorporation efficiency of MDMs in spheroids

Monotypic and heterotypic spheroids were used to recruit monocytes and macrophages as before using prestained 5000 pre-stained Monocytes/macrophages (CellTracker™ green). After 16-18hrs, the collagen matrix was dissolved by adding acidified medium to the well. Spheroids were harvested using wide bore tips and were allowed to settle by gravity. The harvested spheroids were washed twice-thrice with PBS to remove unincorporated and loosely bound monocytes/ macrophages. These spheroids were then dissociated using Accutase and neutralized by adding equal volume of complete medium. The single cell suspension was centrifuged at 1200x g at room temperature for 5mins. The cellular pellet was washed thrice with PBS and resuspended in 200µl of PBS. The single cell suspension was analyzed through LSRFortessa™ Cell Analyzer (BD). The Analysis of the recruited monocytes/macrophages was done using FlowJo software (Trial version). The plots of a representative experiments are presented. Average values from multiple experiments were plotted using Origin (Student version).

#### Profiling of chemokines secreted by the 3D spheroids

Monotypic and heterotypic spheroids were fabricated as described earlier. For chemokine profiling, 10 well-formed spheroids were collected and transferred to 10 different wells of 96-well plate and incubated with serum free media for 24hrs. Conditioned media was collected for secretome analysis using the HU Proinflam. Chemokine Panel 1 (13-plex) (Biolegend) following manufacturer’s protocol with minor modifications. Briefly the conditioned media, diluted with assay buffer followed by addition of 12.5μl of capture beads and incubated at 8°C for 18hr with shaking. Beads were separated by centrifugation for 5mins at 300rcf and washed once with washing buffer followed by resuspension and incubation with 12.5μl of biotinylated detection antibodies for 1hr at room temperature with shaking. Finally, 12.5μl of detection reagent (streptavidin conjugated with PE) is added to the mixture and incubated for 30mins at room temperature with shaking. Following separation of beads and wash, the beads were resuspended in 200μl of wash buffer and readings were obtained on a flow cytometer with appropriate assay settings. The fcs files were analyzed using BioLegend’s LEGENDplex^TM^ data analysis software (https://legendplex.qognit.com). The MFI values of samples were used to quantify the analytes using standard graph of respective analyte.

#### Expression profiling of chemokine receptors on activated and differentiated macrophages

Receptors for relevant chemokines were identified and primers were designed (SI Table 5). GAPDH was used as internal control. Monocytes and differentiated subtypes (M0, M1 and M2) were collected and subjected to isolation of total RNA using Trizol. Total RNA (3-5μg) was used as template for cDNA synthesis using RevertAid cDNA synthesis kit. cDNA (50μg) was used as template in each PCR reaction. PCR were done using 2X SYBR green PCR master mix in CFX96 Touch System (Bio-Rad). Ct values were used to calculate the fold change. Mean of fold change values are plotted with standard error.

##### Correlation analysis of CR expression and Macrophage infiltration

Correlation between the expression level of Chemokine receptors (CXCR1, CCR3 and CCR6) and macrophage infiltration efficiency into breast tumor was analyzed using the TIMER2.0 (http://timer.cistrome.org/) database. The correlation analysis was done using TIMER algorithm. The results displayed the purity-adjusted correlation coefficient (spearman’s rho) values.

#### Analysis of impact of recruited activated macrophages (M0) on spheroid behaviour

As MCF7 spheroids exhibit maximum incorporation of M0 macrophages, viability and drug resistance experiments were done with it only.

##### Spheroid viability

MCF7 spheroids were subjected to the incorporation of M0 macrophages for 16-24 hr as before. Control and M0-incoroprated spheroids were retrieved from the collagen matrix and maintained in suspension at standard cell culture conditions for 72 hr. Control and macrophage incorporated spheroids were stained with Calcein AM (1µM for 30mins) and Propidium Iodide (PI, 1 µg/mL for 10mins) at room temperature. The spheroids were washed thrice and imaged using a CLSM (Leica TC SP8) under a 20X objective, with live cells marked by green fluorescence (Calcein AM) and dead cells by red fluorescence (PI). Cell death was analyzed by measuring the mean fluorescent intensity of PI/total spheroid area. Average values from multiple experiments were plotted using Origin.

##### Drug resistance of M0 incorporated spheroid

The control and macrophage incorporated spheroids were retrieved from the collagen and maintained in suspension for 24hrs. Later, control and macrophage incorporated spheroids were treated with Doxorubicin (50 µM) for 72 hours under standard cell culture condition. Post-treatment, cell viability was assessed Calcein-PI staining. MFI of PI/area of spheroids were used to check cell death as before. Average values from multiple experiments were plotted using Origin.

##### U937 and Human monocyte derived activated macrophage (M0) influence spheroids migration

To analyze the impact of incorporated macrophages (M0) on migration ability, spheroids with different composition and migrational ability used. Their migration was analyzed after incorporating macrophages from U937 cell line and healthy human PBMCs. The hMDMs were obtained by treating hMonocytes with 10ng/ml M-CSF for 10-15 days. The differentiated macrophages were further used for recruitment and incorporation in spheroids as before. Monotypic and heterotypic spheroids of mCherry-MCF7 and EGFP-MDA MB231 were fabricated as before. These spheroids are then entrapped with Rat Tail collagen 1 followed by its solidification. Upon polymerization of collagen, 10,000 monocytes/ macrophages were stained with CellTracker Blue (Thermo Fisher Inc) are seeded on the top of collagen gel and incubated under standard culture conditions. After 24 and 72 hrs., the spheroids were analyzed under 10X objective of CLSM (Leica TC SP8). Optical sections of 0.5µ were acquired using Z scanning mode. Post incorporation, invasion of the cells in the surrounding ECM was observed for 24hrs and 72hrs and the images were obtained under confocal microscope. Here spheroids without any incorporated macrophages were used as control.

#### Development of TEM and analysis of TEM-CM on spheroid migration

##### Development of TEM

Naïve (M0) macrophages were differentiated from U937 cells as explained previously in section 2.2.3. Naïve macrophages were then treated with 50% conditioned medium obtained from spheroids for 48hrs. Macrophages treated with MCF-7 spheroid conditioned medium were labeled as Tumor Educated Macrophage-1 (TEM-1). Whereas, those treated with MDA-MB-231 spheroids were called as Tumor Educated Macrophage-2 (TEM-2). Upon polarization into TEMs, the media was discarded and macrophages were washed once with PBS and maintained in 3ml serum free DMEM for overnight. The media was then collected in sterile centrifuge tube and subjected to centrifugation at maximum speed for 5mins at room temperature to remove cellular debris. The supernatant was then aliquoted and used or stored at −20°C until further use.

##### Analyzing the effect of TEM-CM on Spheroid Migration

Well-formed spheroids were harvested using wide bore tips and washed with PBS. Each well of a fresh 96 well plate was coated with 30μl of 1mg/ml collagen-1. The matrix was allowed to polymerized. Upon polymerization spheroids were deposited on the polymerized matrix. The spheroids were then overlaid with 50μl of collagen or Matrigel and allowed it to polymerize. Upon polymerization the matrix was then overlaid with 100μl of conditioned media of TEMs. Here serum free DMEM was used as control. Plates were then incubated at standard tissue culture conditions and imaged at intervals till 72hrs. Radius of the spheroid and the perimeter of the polygon connecting the migrating fronts were measured using MagVision software. Maximum migrating distance was calculated for MCF-7 spheroids and Average of maximum migrating distance was calculated for MDA-MB-231 spheroid to assess their invasiveness.

#### Evaluation of TAM marker (Siglec-1) in monocytes, activated macrophages, and spheroid incorporated macrophages

Total RNA from monocytes, activated and polarized macrophages were isolated as before and used for cDNA synthesis using RevertAid cDNA synthesis kit. Expression of Siglec-1 gene was checked using qRT-PCR as before. Similarly, control spheroids and M0-incorporated spheroids were also subjected to RNA isolation, cDNA synthesis and RT-PCR for Siglec-1 mRNA. Foldchange values were calculated and plotted using Origin.

Expression of Siglec-1 protein in patients’ samples was analyzed by immunohistochemistry. A small piece of tumor sample was immediately transferred to 10%PFA. The sample were then dehydrated with increasing concentration of ethanol and finally with xylene. The dehydrated samples were then subjected to paraffin block fabrication. The blocks were then sectioned and the sections were deparaffinized and rehydrated. The slides were then subjected to antigen retrieval followed by the incubation with anti Siglec-1 antibody (1:500) for overnight at 4°C. The sections were incubated with HRP conjugated secondary antibody and TMB/H_2_O_2_ developing solution. Nuclei were counterstained with hematoxylin. The sections were observed and imaged under the microscope.

##### Correlation analysis of Siglec-1 expression and Macrophage infiltration

Correlation between Siglec1 expression and macrophage incorporation into breast tumor was analyzed using the TIMER2.0 (http://timer.cistrome.org/) database as before. Briefly, the TIMER algorithm-based analysis was used to assess the Siglec-1 expression and macrophage incorporation in different subtypes of breast tumor. The results displayed the purity-adjusted correlation coefficient (spearman’s rho) across the subtypes.

#### Statistical analysis

All experiments were repeated thrice with three technical replicates. Values are plotted as Mean ± SEM. All statistical analysis was performed using OriginPro Software. We have used One-Way ANOVA followed by post hoc analysis (Tukey’s test) for comparison between more than two experimental groups. Additional statistical information has also been provided in figure legends. *P* ≤ 0.05 was considered as statistically significant for all experiments, and values were assigned accordingly (\**P* ≤ 0.05, \*\**P* ≤ 0.005, \*\*\**P* ≤ 0.001).

## Acknowledgement

TD acknowledges funding support from SERB POWER grant along with DST, DBT and ICMR. SSK thanks Swarna Jayanti Fellowship (to S.S.K.) from the Science and Engineering Research Board (SERB), Department of Science and Technology, Government of India (grant number: SB/SJF/2021–22/01). AT thanks DBT for fellowship. RNI thanks SERB-POWER for fellowship. RG thanks ICMR for Junior Research fellowship. SG thanks CSIR for funding.

