## Supplemental Table for "Breast cancer spheroids prefer activated macrophages as an accomplice: An in vitro study"

SI Table 1

Upregulated protein listing in activated macrophage (M0), M1 and M2 population

| M0 vs Mono | M0 vs M1 | M2 vs M2 |
| --- | --- | --- |
| 14-3-3 protein epsilon [Homo sapiens] | 14-3-3 protein theta [Homo sapiens] | 26S proteasome non-ATPase regulatory subunit 1 isoform 2 [Homo sapiens] |
| 14-3-3 protein gamma [Homo sapiens] | 14-3-3 protein zeta/delta isoform X1 [Homo sapiens] | 40S ribosomal protein S13 [Homo sapiens] |
| 14-3-3 protein theta [Homo sapiens] | 40S ribosomal protein S26 [Homo sapiens] | 40S ribosomal protein S2 [Homo sapiens] |
| 14-3-3 protein zeta/delta isoform X1 [Homo sapiens] | 40S ribosomal protein S3 isoform 1 [Homo sapiens] | 40S ribosomal protein S26 [Homo sapiens] |
| 40S ribosomal protein S11 [Homo sapiens] | 60 kDa heat shock protein, mitochondrial [Homo sapiens] | 40S ribosomal protein S3 isoform 1 [Homo sapiens] |
| 40S ribosomal protein S12 [Homo sapiens] | 60S acidic ribosomal protein P2 [Homo sapiens] | 40S ribosomal protein S4, X isoform [Homo sapiens] |
| 40S ribosomal protein S13 [Homo sapiens] | 60S ribosomal protein L13 isoform 1 [Homo sapiens] | 40S ribosomal protein S9 isoform a [Homo sapiens] |
| 40S ribosomal protein S16 isoform 1 [Homo sapiens] | 60S ribosomal protein L18a [Homo sapiens] | 40S ribosomal protein SA isoform 1 [Homo sapiens] |
| 40S ribosomal protein S3 isoform 1 [Homo sapiens] | 60S ribosomal protein L30 [Homo sapiens] | 5'-3' exonuclease PLD3 isoform X1 [Homo sapiens] |
| 40S ribosomal protein S3a isoform 1 [Homo sapiens] | 60S ribosomal protein L7 isoform 1 [Homo sapiens] | 60 kDa heat shock protein, mitochondrial [Homo sapiens] |
| 40S ribosomal protein S4, X isoform [Homo sapiens] | actin-related protein 2/3 complex subunit 2 [Homo sapiens] | 60S acidic ribosomal protein P0 [Homo sapiens] |
| 40S ribosomal protein SA isoform 1 [Homo sapiens] | actin-related protein 2/3 complex subunit 4 isoform a [Homo sapiens] | 60S ribosomal protein L10 isoform a [Homo sapiens] |
| 60S ribosomal protein L12 [Homo sapiens] | actin-related protein 3 isoform 1 [Homo sapiens] | 60S ribosomal protein L12 [Homo sapiens] |
| 60S ribosomal protein L7 isoform 1 [Homo sapiens] | adenylyl cyclase-associated protein 1 isoform X1 [Homo sapiens] | 60S ribosomal protein L13 isoform 1 [Homo sapiens] |
| 60S ribosomal protein L7a [Homo sapiens] | annexin A1 isoform X2 [Homo sapiens] | 60S ribosomal protein L18 isoform 1 [Homo sapiens] |
| 6-phosphogluconate dehydrogenase, decarboxylating isoform 3 [Homo sapiens] | annexin A6 isoform 1 [Homo sapiens] | 60S ribosomal protein L24 [Homo sapiens] |
| 7-dehydrocholesterol reductase [Homo sapiens] | ATP-dependent RNA helicase DDX3X isoform 1 [Homo sapiens] | 60S ribosomal protein L27 [Homo sapiens] |
| acetyl-CoA acetyltransferase, cytosolic isoform 1 [Homo sapiens] | calnexin isoform b [Homo sapiens] | 60S ribosomal protein L3 isoform b [Homo sapiens] |
| acid ceramidase isoform X1 [Homo sapiens] | calreticulin precursor [Homo sapiens] | 60S ribosomal protein L30 [Homo sapiens] |
| actin, cytoplasmic 1 [Homo sapiens] | cystatin-B [Homo sapiens] | 60S ribosomal protein L7 isoform 1 [Homo sapiens] |
| actin-related protein 2/3 complex subunit 2 [Homo sapiens] | DNA-dependent protein kinase catalytic subunit isoform 1 [Homo sapiens] | 60S ribosomal protein L7a [Homo sapiens] |
| actin-related protein 3 isoform 1 [Homo sapiens] | endoplasmic reticulum chaperone BiP precursor [Homo sapiens] | 60S ribosomal protein L9 [Homo sapiens] |
| activated RNA polymerase II transcriptional coactivator p15 isoform X1 [Homo sapiens] | eukaryotic initiation factor 4A-I isoform 1 [Homo sapiens] | acetyl-CoA acetyltransferase, mitochondrial isoform b precursor [Homo sapiens] |
| adenosylhomocysteinase isoform 1 [Homo sapiens] | fatty acid-binding protein 5 [Homo sapiens] | acid ceramidase isoform X1 [Homo sapiens] |
| adenylate kinase 2, mitochondrial isoform f [Homo sapiens] | glucose-6-phosphate 1-dehydrogenase isoform b [Homo sapiens] | actin-related protein 3 isoform 1 [Homo sapiens] |
| adenylyl cyclase-associated protein 1 isoform X1 [Homo sapiens] | glyceraldehyde-3-phosphate dehydrogenase isoform 1 [Homo sapiens] | adenine phosphoribosyltransferase isoform a [Homo sapiens] |
| alpha-actinin-4 isoform X2 [Homo sapiens] | heterogeneous nuclear ribonucleoprotein A3 isoform a [Homo sapiens] | adenylate kinase 2, mitochondrial isoform f [Homo sapiens] |
| alpha-enolase isoform 1 [Homo sapiens] | HLA class I histocompatibility antigen, A alpha chain A*03:01:01:01 precursor [Homo sapiens] | ADP-ribosylation factor-like protein 8B [Homo sapiens] |
| aminopeptidase B isoform a [Homo sapiens] | hypoxia up-regulated protein 1 precursor [Homo sapiens] | alkyldihydroxyacetonephosphate synthase, peroxisomal precursor [Homo sapiens] |
| annexin A2 isoform 1 [Homo sapiens] | intercellular adhesion molecule 1 precursor [Homo sapiens] | alpha-1,3-mannosyl-glycoprotein 2-beta-N-acetylglucosaminyltransferase [Homo sapiens] |
| annexin A5 [Homo sapiens] | interstitial collagenase isoform 1 preproprotein [Homo sapiens] | alpha-galactosidase A precursor [Homo sapiens] |
| annexin A6 isoform 1 [Homo sapiens] | keratin, type I cytoskeletal 10 isoform 1 [Homo sapiens] | aminopeptidase N [Homo sapiens] |
| apoptosis-inducing factor 1, mitochondrial isoform AIF precursor [Homo sapiens] | keratin, type II cytoskeletal 1 [Homo sapiens] | annexin A1 isoform X1 [Homo sapiens] |
| ATP-dependent 6-phosphofructokinase, liver type isoform X3 [Homo sapiens] | lamin isoform A [Homo sapiens] | annexin A11 isoform X3 [Homo sapiens] |
| beta-galactosidase isoform d precursor [Homo sapiens] | liver carboxylesterase 1 isoform a precursor [Homo sapiens] | annexin A2 isoform 1 [Homo sapiens] |
| bifunctional purine biosynthesis protein ATIC [Homo sapiens] | long-chain-fatty-acid--CoA ligase 1 isoform a [Homo sapiens] | annexin A4 isoform a [Homo sapiens] |
| bolA-like protein 2 isoform 1 [Homo sapiens] | metalloproteinase inhibitor 1 precursor [Homo sapiens] | annexin A6 isoform 1 [Homo sapiens] |
| C-1-tetrahydrofolate synthase, cytoplasmic isoform 1 [Homo sapiens] | NADPH--cytochrome P450 reductase isoform 3 [Homo sapiens] | ATP synthase subunit alpha, mitochondrial isoform a precursor [Homo sapiens] |
| CAD protein isoform 2 [Homo sapiens] | neutral alpha-glucosidase AB isoform 3 precursor [Homo sapiens] | ATP-dependent RNA helicase DDX1 [Homo sapiens] |
| calnexin isoform X2 [Homo sapiens] | OCIA domain-containing protein 1 isoform 1 [Homo sapiens] | beta-hexosaminidase subunit alpha isoform 2 preproprotein [Homo sapiens] |
| catalase [Homo sapiens] | phosphoglycerate kinase 1 [Homo sapiens] | beta-hexosaminidase subunit beta isoform 1 preproprotein [Homo sapiens] |
| catechol O-methyltransferase isoform S-COMT [Homo sapiens] | phosphoglycerate mutase 1 isoform 1 [Homo sapiens] | calreticulin precursor [Homo sapiens] |
| cathepsin D preproprotein [Homo sapiens] | plasminogen activator inhibitor 2 [Homo sapiens] | catalase [Homo sapiens] |
| cathepsin G isoform X1 [Homo sapiens] | plastin-2 isoform X1 [Homo sapiens] | catechol O-methyltransferase isoform MB-COMT [Homo sapiens] |
| chitinase-3-like protein 1 precursor [Homo sapiens] | plectin isoform X3 [Homo sapiens] | cathepsin S isoform 2 preproprotein [Homo sapiens] |
| chloride intracellular channel protein 1 [Homo sapiens] | prohibitin-2 isoform 1 [Homo sapiens] | cell cycle progression protein 1 isoform 1 [Homo sapiens] |
| clathrin heavy chain 1 isoform 2 [Homo sapiens] | proteasome subunit alpha type-7 [Homo sapiens] | clathrin heavy chain 1 isoform 2 [Homo sapiens] |
| core histone macro-H2A.1 isoform 2 [Homo sapiens] | protein disulfide-isomerase A3 precursor [Homo sapiens] | cytochrome b-c1 complex subunit 1, mitochondrial precursor [Homo sapiens] |
| coronin-1A isoform X2 [Homo sapiens] | protein disulfide-isomerase A4 isofrom 2 precursor [Homo sapiens] | cytochrome c oxidase subunit 5A, mitochondrial precursor [Homo sapiens] |
| CTP synthase 1 isoform a [Homo sapiens] | pyruvate kinase PKM isoform X4 [Homo sapiens] | dihydrolipoyl dehydrogenase, mitochondrial isoform 1 precursor [Homo sapiens] |
| cullin-associated NEDD8-dissociated protein 1 isoform 1 [Homo sapiens] | receptor of activated protein C kinase 1 [Homo sapiens] | DNA topoisomerase 1 [Homo sapiens] |
| cytochrome b-c1 complex subunit 2, mitochondrial precursor [Homo sapiens] | rRNA 2'-O-methyltransferase fibrillarin isoform X1 [Homo sapiens] | DNA-3-methyladenine glycosylase isoform c [Homo sapiens] |
| cytochrome c [Homo sapiens] | splicing factor, proline- and glutamine-rich isoform X1 [Homo sapiens] | DNA-dependent protein kinase catalytic subunit isoform 1 [Homo sapiens] |
| cytosolic acyl coenzyme A thioester hydrolase isoform hBACHa [Homo sapiens] | T-complex protein 1 subunit delta isoform a [Homo sapiens] | dnaJ homolog subfamily B member 11 isoform 1 precursor [Homo sapiens] |
| D-3-phosphoglycerate dehydrogenase isoform X1 [Homo sapiens] | transitional endoplasmic reticulum ATPase isoform 1 [Homo sapiens] | dolichyl-diphosphooligosaccharide--protein glycosyltransferase subunit 1 precursor [Homo sapiens] |
| deoxyuridine 5'-triphosphate nucleotidohydrolase, mitochondrial isoform 1 precursor [Homo sapiens] | transketolase isoform X1 [Homo sapiens] | dolichyl-diphosphooligosaccharide--protein glycosyltransferase subunit 2 isoform 1 precursor [Homo sapiens] |
| dipeptidyl peptidase 1 isoform a preproprotein [Homo sapiens] | ubiquitin-like protein ISG15 [Homo sapiens] | E3 SUMO-protein ligase RanBP2 isoform X6 [Homo sapiens] |
| DNA replication licensing factor MCM2 [Homo sapiens] | vimentin [Homo sapiens] | electron transfer flavoprotein subunit alpha, mitochondrial isoform a [Homo sapiens] |
| DNA replication licensing factor MCM3 isoform 1 [Homo sapiens] | voltage-dependent anion-selective channel protein 1 isoform X1 [Homo sapiens] | elongation factor 2 [Homo sapiens] |
| DNA replication licensing factor MCM7 isoform 1 [Homo sapiens] |  | elongation factor Tu, mitochondrial isoform 1 precursor [Homo sapiens] |
| DNA topoisomerase 1 [Homo sapiens] |  | endoplasmic reticulum chaperone BiP precursor [Homo sapiens] |
| DNA topoisomerase 2-beta isoform X1 [Homo sapiens] |  | endoplasmin precursor [Homo sapiens] |
| DNA-(apurinic or apyrimidinic site) endonuclease [Homo sapiens] |  | ERO1-like protein alpha isoform 2 precursor [Homo sapiens] |
| DNA-dependent protein kinase catalytic subunit isoform 1 [Homo sapiens] |  | exportin-1 isoform X3 [Homo sapiens] |
| dolichyl-diphosphooligosaccharide--protein glycosyltransferase subunit 2 isoform 1 precursor [Homo sapiens] |  | fatty acid synthase isoform X1 [Homo sapiens] |
| E3 ubiquitin-protein ligase HUWE1 isoform X9 [Homo sapiens] |  | fibronectin type III domain-containing protein 3B [Homo sapiens] |
| electron transfer flavoprotein subunit beta isoform 2 [Homo sapiens] |  | filamin-A isoform 2 [Homo sapiens] |
| elongation factor 1-delta isoform X1 [Homo sapiens] |  | fructose-bisphosphate aldolase A isoform 1 [Homo sapiens] |
| elongation factor 1-gamma [Homo sapiens] |  | glucose-6-phosphate isomerase isoform X1 [Homo sapiens] |
| elongation factor 2 [Homo sapiens] |  | glutamate dehydrogenase 1, mitochondrial isoform a precursor [Homo sapiens] |
| endoplasmic reticulum chaperone BiP precursor [Homo sapiens] |  | glutaminase kidney isoform, mitochondrial isoform 2 [Homo sapiens] |
| endoplasmin precursor [Homo sapiens] |  | glyceraldehyde-3-phosphate dehydrogenase isoform 1 [Homo sapiens] |
| erlin-1 isoform a [Homo sapiens] |  | glycogen phosphorylase, liver form isoform 1 [Homo sapiens] |
| eukaryotic translation initiation factor 2 subunit 3 [Homo sapiens] |  | heat shock 70 kDa protein 1A [Homo sapiens] |
| eukaryotic translation initiation factor 3 subunit A [Homo sapiens] |  | heat shock cognate 71 kDa protein isoform X1 [Homo sapiens] |
| eukaryotic translation initiation factor 3 subunit B isoform 1 [Homo sapiens] |  | heterogeneous nuclear ribonucleoprotein A1 isoform a [Homo sapiens] |
| eukaryotic translation initiation factor 3 subunit L isoform 1 [Homo sapiens] |  | heterogeneous nuclear ribonucleoprotein A3 isoform a [Homo sapiens] |
| eukaryotic translation initiation factor 6 isoform a [Homo sapiens] |  | heterogeneous nuclear ribonucleoprotein D0 isoform d [Homo sapiens] |
| exportin-1 isoform X3 [Homo sapiens] |  | heterogeneous nuclear ribonucleoprotein F [Homo sapiens] |
| F-actin-capping protein subunit alpha-1 isoform X2 [Homo sapiens] |  | heterogeneous nuclear ribonucleoprotein H3 isoform b [Homo sapiens] |
| far upstream element-binding protein 2 isoform 1 [Homo sapiens] |  | heterogeneous nuclear ribonucleoprotein K isoform d [Homo sapiens] |
| fatty acid synthase isoform X1 [Homo sapiens] |  | heterogeneous nuclear ribonucleoprotein M isoform X4 [Homo sapiens] |
| fatty acid-binding protein 5 [Homo sapiens] |  | heterogeneous nuclear ribonucleoprotein R isoform X2 [Homo sapiens] |
| fermitin family homolog 3 long isoform [Homo sapiens] |  | heterogeneous nuclear ribonucleoprotein U isoform a [Homo sapiens] |
| fructose-bisphosphate aldolase A isoform 1 [Homo sapiens] |  | heterogeneous nuclear ribonucleoprotein U-like protein 1 isoform e [Homo sapiens] |
| galectin-3 isoform 3 [Homo sapiens] |  | heterogeneous nuclear ribonucleoproteins A2/B1 isoform X1 [Homo sapiens] |
| glucose-6-phosphate isomerase isoform X1 [Homo sapiens] |  | histone-binding protein RBBP4 isoform a [Homo sapiens] |
| glutamate dehydrogenase 1, mitochondrial isoform a precursor [Homo sapiens] |  | hydroxyacyl-coenzyme A dehydrogenase, mitochondrial isoform 3 [Homo sapiens] |
| glutathione reductase, mitochondrial isoform 1 precursor [Homo sapiens] |  | interleukin enhancer-binding factor 2 isoform 1 [Homo sapiens] |
| glutathione S-transferase omega-1 isoform 1 [Homo sapiens] |  | interleukin enhancer-binding factor 3 isoform a [Homo sapiens] |
| glutathione S-transferase P [Homo sapiens] |  | leucine-rich PPR motif-containing protein, mitochondrial isoform X1 [Homo sapiens] |
| glyceraldehyde-3-phosphate dehydrogenase isoform 1 [Homo sapiens] |  | lupus La protein [Homo sapiens] |
| GTP-binding nuclear protein Ran isoform 1 [Homo sapiens] |  | lysosomal alpha-mannosidase isoform X1 [Homo sapiens] |
| guanine nucleotide-binding protein G(I)/G(S)/G(T) subunit beta-1 isoform 1 [Homo sapiens] |  | lysosome-associated membrane glycoprotein 1 isoform X1 [Homo sapiens] |
| heat shock cognate 71 kDa protein isoform X1 [Homo sapiens] |  | lysosome-associated membrane glycoprotein 2 isoform B precursor [Homo sapiens] |
| heat shock protein beta-1 [Homo sapiens] |  | macrophage-capping protein isoform X1 [Homo sapiens] |
| heat shock protein HSP 90-alpha isoform X1 [Homo sapiens] |  | malate dehydrogenase, mitochondrial isoform 1 precursor [Homo sapiens] |
| heat shock protein HSP 90-beta isoform c [Homo sapiens] |  | matrin-3 isoform a [Homo sapiens] |
| heterochromatin protein 1-binding protein 3 isoform 1 [Homo sapiens] |  | matrix metalloproteinase-9 preproprotein [Homo sapiens] |
| heterogeneous nuclear ribonucleoprotein F [Homo sapiens] |  | microsomal glutathione S-transferase 3 isoform X1 [Homo sapiens] |
| heterogeneous nuclear ribonucleoprotein K isoform b [Homo sapiens] |  | myeloid-derived growth factor precursor [Homo sapiens] |
| heterogeneous nuclear ribonucleoprotein M isoform X4 [Homo sapiens] |  | NADPH--cytochrome P450 reductase isoform 1 [Homo sapiens] |
| heterogeneous nuclear ribonucleoprotein U isoform a [Homo sapiens] |  | non-POU domain-containing octamer-binding protein isoform 1 [Homo sapiens] |
| heterogeneous nuclear ribonucleoprotein U-like protein 2 [Homo sapiens] |  | NPC intracellular cholesterol transporter 1 isoform X5 [Homo sapiens] |
| high mobility group protein B2 [Homo sapiens] |  | nucleolar RNA helicase 2 isoform 1 [Homo sapiens] |
| histone deacetylase 1 [Homo sapiens] |  | osteopontin isoform OPN-a precursor [Homo sapiens] |
| HLA class I histocompatibility antigen, C alpha chain precursor [Homo sapiens] |  | palmitoyl-protein thioesterase 1 isoform 1 precursor [Homo sapiens] |
| hsc70-interacting protein isoform 2 [Homo sapiens] |  | peptidyl-prolyl cis-trans isomerase B precursor [Homo sapiens] |
| hypoxanthine-guanine phosphoribosyltransferase [Homo sapiens] |  | peptidyl-prolyl cis-trans isomerase FKBP1A isoform a [Homo sapiens] |
| hypoxia up-regulated protein 1 precursor [Homo sapiens] |  | peroxiredoxin-4 precursor [Homo sapiens] |
| importin subunit beta-1 isoform 1 [Homo sapiens] |  | peroxisomal multifunctional enzyme type 2 isoform 2 [Homo sapiens] |
| importin-5 isoform X1 [Homo sapiens] |  | phosphate carrier protein, mitochondrial isoform b precursor [Homo sapiens] |
| inositol 1,4,5-trisphosphate receptor type 1 isoform X13 [Homo sapiens] |  | plectin isoform X3 [Homo sapiens] |
| integrin beta-2 isoform 2 [Homo sapiens] |  | polypyrimidine tract-binding protein 1 isoform X2 [Homo sapiens] |
| interstitial collagenase isoform 1 preproprotein [Homo sapiens] |  | pre-mRNA-processing factor 19 [Homo sapiens] |
| lactoylglutathione lyase [Homo sapiens] |  | pre-mRNA-processing-splicing factor 8 [Homo sapiens] |
| lamin isoform A [Homo sapiens] |  | presequence protease, mitochondrial isoform 2 precursor [Homo sapiens] |
| lamin-B1 isoform 1 [Homo sapiens] |  | profilin-1 isoform 2 [Homo sapiens] |
| leucine-rich PPR motif-containing protein, mitochondrial isoform X2 [Homo sapiens] |  | prohibitin isoform 1 [Homo sapiens] |
| leukocyte elastase inhibitor isoform X2 [Homo sapiens] |  | proteasome subunit alpha type-3 isoform 1 [Homo sapiens] |
| liver carboxylesterase 1 isoform b precursor [Homo sapiens] |  | proteasome subunit alpha type-4 isoform 1 [Homo sapiens] |
| L-lactate dehydrogenase A chain isoform 1 [Homo sapiens] |  | proteasome subunit alpha type-6 isoform a [Homo sapiens] |
| long-chain-fatty-acid--CoA ligase 1 isoform c [Homo sapiens] |  | protein disulfide-isomerase A3 precursor [Homo sapiens] |
| long-chain-fatty-acid--CoA ligase 4 isoform 1 [Homo sapiens] |  | protein disulfide-isomerase A4 isofrom 2 precursor [Homo sapiens] |
| lupus La protein [Homo sapiens] |  | protein disulfide-isomerase precursor [Homo sapiens] |
| lysosome-associated membrane glycoprotein 2 isoform B precursor [Homo sapiens] |  | protein PML isoform 1 [Homo sapiens] |
| macrophage-capping protein isoform X1 [Homo sapiens] |  | pyruvate kinase PKM isoform X4 [Homo sapiens] |
| malate dehydrogenase, peroxisomal isoform MDH1x [Homo sapiens] |  | receptor of activated protein C kinase 1 [Homo sapiens] |
| matrix metalloproteinase-9 preproprotein [Homo sapiens] |  | retinol dehydrogenase 11 isoform 1 precursor [Homo sapiens] |
| medium-chain specific acyl-CoA dehydrogenase, mitochondrial isoform e [Homo sapiens] |  | ribonuclease inhibitor isoform X1 [Homo sapiens] |
| microsomal glutathione S-transferase 1 isoform a [Homo sapiens] |  | ribosome-binding protein 1 isoform 2 [Homo sapiens] |
| mitochondrial carrier homolog 1 isoform PSAP-LS [Homo sapiens] |  | RNA-binding motif protein, X chromosome isoform 1 [Homo sapiens] |
| moesin isoform X1 [Homo sapiens] |  | RNA-binding protein 25 isoform X1 [Homo sapiens] |
| myosin light polypeptide 6 isoform 2 [Homo sapiens] |  | RNA-binding protein 39 isoform b [Homo sapiens] |
| myosin-9 [Homo sapiens] |  | rRNA 2'-O-methyltransferase fibrillarin isoform X2 [Homo sapiens] |
| NME1-NME2 protein [Homo sapiens] |  | RRP12-like protein isoform 3 [Homo sapiens] |
| NPC intracellular cholesterol transporter 1 isoform X5 [Homo sapiens] |  | sarcoplasmic/endoplasmic reticulum calcium ATPase 2 isoform X1 [Homo sapiens] |
| nuclear pore membrane glycoprotein 210 precursor [Homo sapiens] |  | serine beta-lactamase-like protein LACTB, mitochondrial isoform a precursor [Homo sapiens] |
| nucleolar RNA helicase 2 isoform 1 [Homo sapiens] |  | serine/arginine-rich splicing factor 1 isoform 1 [Homo sapiens] |
| obg-like ATPase 1 isoform 1 [Homo sapiens] |  | signal recognition particle receptor subunit alpha isoform 2 [Homo sapiens] |
| peptidyl-prolyl cis-trans isomerase A isoform 1 [Homo sapiens] |  | small nuclear ribonucleoprotein Sm D3 [Homo sapiens] |
| peptidyl-prolyl cis-trans isomerase B precursor [Homo sapiens] |  | splicing factor 3A subunit 1 [Homo sapiens] |
| peptidyl-prolyl cis-trans isomerase FKBP4 isoform X1 [Homo sapiens] |  | splicing factor 3B subunit 1 isoform 1 [Homo sapiens] |
| peptidyl-prolyl cis-trans isomerase FKBP5 isoform 1 [Homo sapiens] |  | splicing factor 3B subunit 3 [Homo sapiens] |
| peroxiredoxin-1 [Homo sapiens] |  | splicing factor, proline- and glutamine-rich isoform X1 [Homo sapiens] |
| peroxiredoxin-6 [Homo sapiens] |  | squalene monooxygenase isoform X1 [Homo sapiens] |
| peroxisomal multifunctional enzyme type 2 isoform 2 [Homo sapiens] |  | staphylococcal nuclease domain-containing protein 1 [Homo sapiens] |
| persulfide dioxygenase ETHE1, mitochondrial isoform 1 [Homo sapiens] |  | stress-70 protein, mitochondrial precursor [Homo sapiens] |
| phosphate carrier protein, mitochondrial isoform b precursor [Homo sapiens] |  | T-complex protein 1 subunit zeta isoform a [Homo sapiens] |
| phosphoglycerate kinase 1 [Homo sapiens] |  | three-prime repair exonuclease 1 isoform c [Homo sapiens] |
| phosphoglycerate mutase 1 isoform 1 [Homo sapiens] |  | transformer-2 protein homolog beta isoform X1 [Homo sapiens] |
| plasminogen activator inhibitor 2 [Homo sapiens] |  | transitional endoplasmic reticulum ATPase isoform 1 [Homo sapiens] |
| plastin-2 isoform X1 [Homo sapiens] |  | transmembrane protein 33 isoform X1 [Homo sapiens] |
| pleckstrin [Homo sapiens] |  | trifunctional enzyme subunit alpha, mitochondrial precursor [Homo sapiens] |
| plectin isoform X11 [Homo sapiens] |  | trifunctional purine biosynthetic protein adenosine-3 isoform X1 [Homo sapiens] |
| poly(rC)-binding protein 1 [Homo sapiens] |  | tripeptidyl-peptidase 1 preproprotein [Homo sapiens] |
| poly(rC)-binding protein 2 isoform e [Homo sapiens] |  | U2 small nuclear ribonucleoprotein A' [Homo sapiens] |
| polypyrimidine tract-binding protein 1 isoform X2 [Homo sapiens] |  | ubiquitin-40S ribosomal protein S27a precursor [Homo sapiens] |
| polyribonucleotide nucleotidyltransferase 1, mitochondrial precursor [Homo sapiens] |  | UDP-glucose:glycoprotein glucosyltransferase 1 precursor [Homo sapiens] |
| probable ATP-dependent RNA helicase DDX17 isoform 1 [Homo sapiens] |  | very long-chain specific acyl-CoA dehydrogenase, mitochondrial isoform 1 precursor [Homo sapiens] |
| proliferation-associated protein 2G4 [Homo sapiens] |  | vimentin [Homo sapiens] |
| proteasome subunit alpha type-5 isoform 1 [Homo sapiens] |  | voltage-dependent anion-selective channel protein 1 isoform X1 [Homo sapiens] |
| proteasome subunit alpha type-6 isoform a [Homo sapiens] |  | V-type proton ATPase 116 kDa subunit a3 isoform a [Homo sapiens] |
| protein disulfide-isomerase A3 precursor [Homo sapiens] |  | V-type proton ATPase subunit B, brain isoform [Homo sapiens] |
| protein disulfide-isomerase A4 isofrom 2 precursor [Homo sapiens] |  | X-ray repair cross-complementing protein 6 isoform 1 [Homo sapiens] |
| protein disulfide-isomerase precursor [Homo sapiens] |  |  |
| pyruvate kinase PKM isoform X4 [Homo sapiens] |  |  |
| rab GDP dissociation inhibitor beta isoform 1 [Homo sapiens] |  |  |
| receptor of activated protein C kinase 1 [Homo sapiens] |  |  |
| retinol dehydrogenase 11 isoform 1 precursor [Homo sapiens] |  |  |
| RNA-binding protein 14 isoform 1 [Homo sapiens] |  |  |
| ruvB-like 1 isoform X3 [Homo sapiens] |  |  |
| ruvB-like 2 isoform 1 [Homo sapiens] |  |  |
| S-formylglutathione hydrolase isoform X1 [Homo sapiens] |  |  |
| stress-70 protein, mitochondrial precursor [Homo sapiens] |  |  |
| sulfide:quinone oxidoreductase, mitochondrial [Homo sapiens] |  |  |
| synaptic vesicle membrane protein VAT-1 homolog [Homo sapiens] |  |  |
| talin-1 [Homo sapiens] |  |  |
| T-complex protein 1 subunit delta isoform a [Homo sapiens] |  |  |
| T-complex protein 1 subunit epsilon isoform e [Homo sapiens] |  |  |
| T-complex protein 1 subunit eta isoform X1 [Homo sapiens] |  |  |
| T-complex protein 1 subunit gamma isoform a [Homo sapiens] |  |  |
| thioredoxin domain-containing protein 5 isoform 1 precursor [Homo sapiens] |  |  |
| thioredoxin isoform 1 [Homo sapiens] |  |  |
| transaldolase [Homo sapiens] |  |  |
| transcription intermediary factor 1-beta [Homo sapiens] |  |  |
| transitional endoplasmic reticulum ATPase isoform 1 [Homo sapiens] |  |  |
| transketolase isoform X1 [Homo sapiens] |  |  |
| transmembrane protein 33 isoform X1 [Homo sapiens] |  |  |
| trifunctional enzyme subunit alpha, mitochondrial precursor [Homo sapiens] |  |  |
| trifunctional purine biosynthetic protein adenosine-3 isoform X1 [Homo sapiens] |  |  |
| triosephosphate isomerase isoform 1 [Homo sapiens] |  |  |
| tubulin alpha-1A chain isoform 2 [Homo sapiens] |  |  |
| tubulin beta chain isoform b [Homo sapiens] |  |  |
| tubulin beta-3 chain isoform 1 [Homo sapiens] |  |  |
| U2 small nuclear ribonucleoprotein A' [Homo sapiens] |  |  |
| ubiquitin-like modifier-activating enzyme 1 isoform X3 [Homo sapiens] |  |  |
| ubiquitin-like protein ISG15 [Homo sapiens] |  |  |
| UPF0687 protein C20orf27 isoform 1 [Homo sapiens] |  |  |
| urokinase plasminogen activator surface receptor isoform 3 precursor [Homo sapiens] |  |  |
| vacuolar protein sorting-associated protein 35 isoform X1 [Homo sapiens] |  |  |
| valine--tRNA ligase [Homo sapiens] |  |  |
| vesicle-trafficking protein SEC22b precursor [Homo sapiens] |  |  |
| vimentin [Homo sapiens] |  |  |
| voltage-dependent anion-selective channel protein 2 isoform 2 [Homo sapiens] |  |  |
| voltage-dependent anion-selective channel protein 3 isoform 1 [Homo sapiens] |  |  |
| V-type proton ATPase catalytic subunit A [Homo sapiens] |  |  |
| X-ray repair cross-complementing protein 5 [Homo sapiens] |  |  |
| X-ray repair cross-complementing protein 6 isoform 1 [Homo sapiens] |  |  |
| Y-box-binding protein 1 [Homo sapiens] |  |  |
| zyxin isoform 1 [Homo sapiens] |  |  |

SI Table 2

LEGENDplexTM Hu Growth factor Panel (13-plex)

| Sr. No. | Name |
| --- | --- |
| 1 | GM-CSF |
| 2 | G-CSF |
| 3 | FGF-Basic |
| 4 | EPO |
| 5 | EGF |
| 6 | Angiopoietin |
| 7 | VEGF |
| 8 | TGF-α |
| 9 | SCF |
| 10 | PDGF-BB |
| 11 | PDGF-AA |
| 12 | M-CSF |
| 13 | HGF |

SI Table 3

LEGENDplex™ HU Essential Immune Response Panel (13-plex)

| Sr. No. | Name |
| --- | --- |
| 1 | IL-4 |
| 2 | IL-2 |
| 3 | CXCL-10 |
| 4 | IL-1β |
| 5 | TNF-α |
| 6 | CCL-2 |
| 7 | IL-17a |
| 8 | IL-6 |
| 9 | IL-10 |
| 10 | IFN-γ |
| 11 | IL-12p70 |
| 12 | CXCL-8 |
| 13 | TGF-β1 |

SI Table 4

LEGENDplex™ Hu Pro-inflam. chemokine panel (13plex)

| Sr. No. | Name |
| --- | --- |
| 1 | CXCL-8 |
| 2 | CXCL-10 |
| 3 | CCL-11 |
| 4 | CCL-17 |
| 5 | CCL-2 |
| 6 | CCL-5 |
| 7 | CCL-3 |
| 8 | CXCL-9 |
| 9 | CXCL-5 |
| 10 | CCL-20 |
| 11 | CXCL-1 |
| 12 | CXCL-11 |
| 13 | CCL-4 |

SI Table 5

Gene name and Primer Seq

| **Sr No.** | **Gene Name** | **Forward Primer** | **Reverse Primer** |
| --- | --- | --- | --- |
| 1 | CD-64 | ATCCACAGAGGCTGGCTACT | ATGCTTTCCCATGCCTGAGC |
| 2 | CD-86 | CCGTCAGTCCTGGCATTATT | ACTCAGTCCCATAGTGCTGT |
| 3 | CD-163 | CCAGGCAACAAACACATGGC | CCACGTGTCACCATGCTTCA |
| 4 | MRC-2 | CATCAACGGCCTCCTCACTG | ATGCTGCAGTCACGGTTCTG |
| 5 | CD-23 | GCCTGTGACGACATGGAAGG | GCCCAGTTGCTGTAGTCCAC |
| 6 | TNF-a | CTGCACTTTGGAGTGATCGG | AGGGTTTGCTACAACATGGG |
| 7 | IL-1b | AGCTGGAGAGTGTAGATCCC | GGACTCTCTGGGTACAGCTC |
| 8 | INF-a-1 | GTCACCCATCTCAGCAAGCC | GTGCCAGGAGCATCAAGGTC |
| 9 | IL-10 | CGGCGCTGTCATCGATTTCT | AGTCGCCACCCTGATGTCTC |
| 10 | CCL-2 | CCGAGAGGCTGAGACTAACC | GGCATTGATTGCATCTGGCT |
| 11 | CCR-2 | GACCGAGTGAGCTCAACATTT | AACCCAACTGAGACTTCTTGC |
| 12 | CXCR-1 | CCATCTCAGGTGTGTTGCAG | ACATGTGTTCCAGCTCCTCA |
| 13 | CXCR-2 | AGCTCTGACTACCACCCAAC | CAGCTGTGACCTGCTGTTAT |
| 14 | CXCR-3 | TGGTGGTCGTGGTGGCCTTT | ACCCTGCTTTCTCGGCCACA |
| 15 | CCR-3 | GGGAGCACACACAAACCATT | CCCTTGGGTGGAACTTGTTG |
| 16 | CCR-5 | CGCTTCTGCAAATGCTGTTC | GGATCGGGTGTAAACTGAGC |
| 17 | CCR-6 | CCGGCTAGTGTTTCCTCTCA | CACCTGTTCTGCCATTGTCC |
| 18 | Siglec-1 | CCTCGGGGAGGAACATCCTT | AGGCGTACCCCATCCTTGA |
