## Supplemental figure for "Breast cancer spheroids prefer activated macrophages as an accomplice: An in vitro study"

SI Fig 1

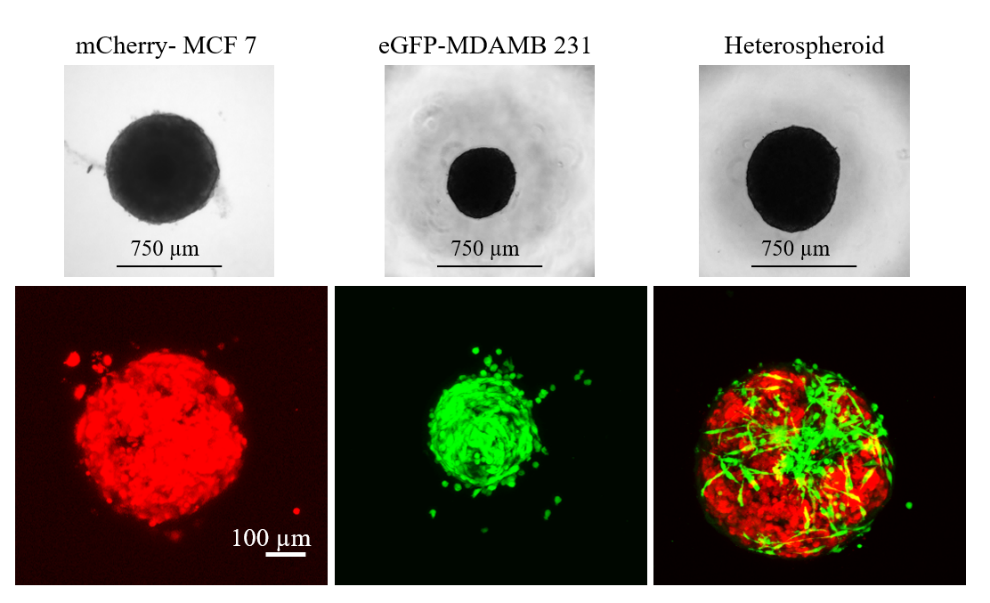

SI Figure 1: Phase contrast (upper panel) and confocal microscopic images (lower panel) of monotypic (MCF7 or MDA MB231) and heterotypic spheroids. Red cells represent mCherry-MCF and green cells represent eGFP-MDA MB231 population.

SI Fig 2

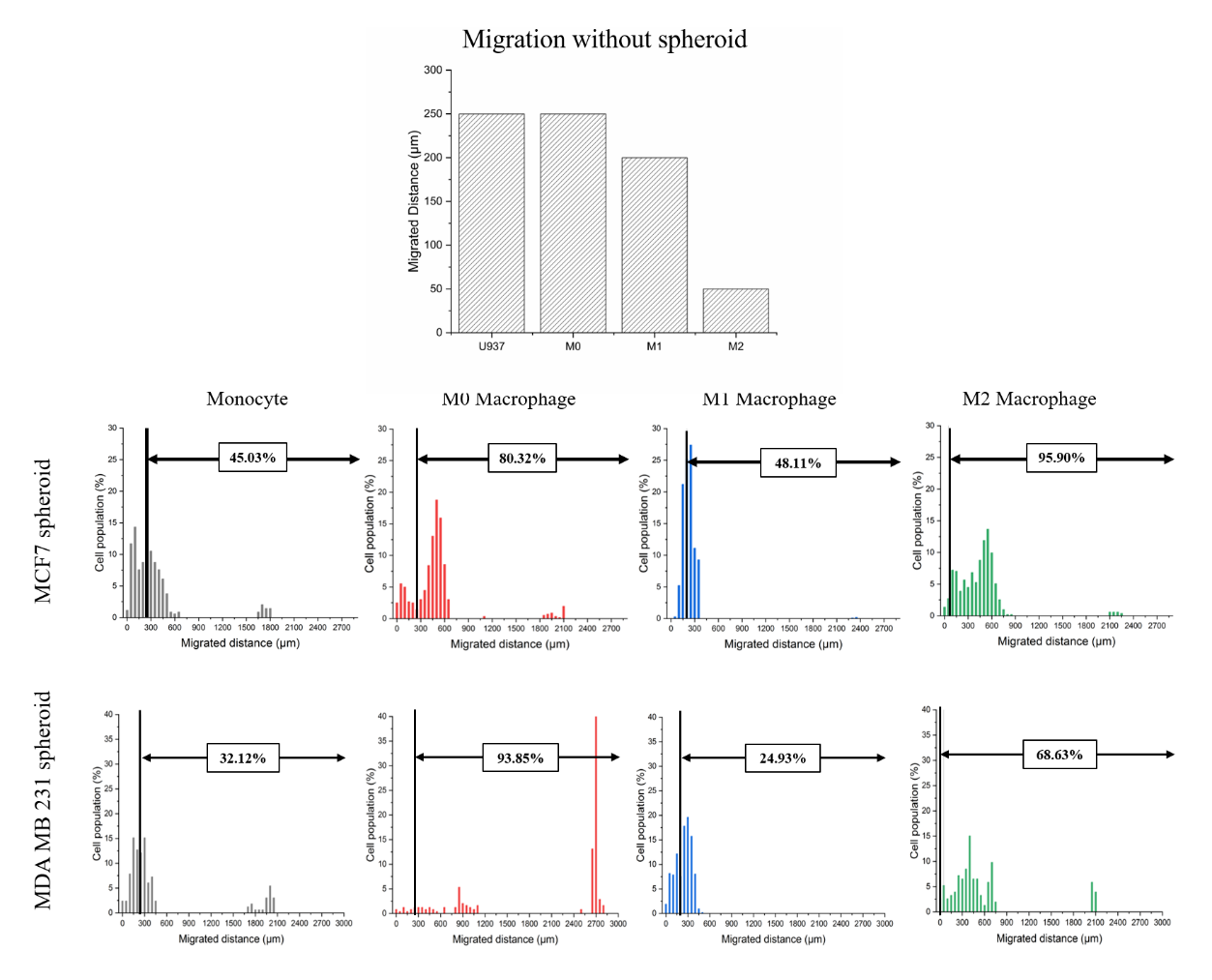

SI Figure 2: Migrated distance by Monocytes, M0, M1 and M2 macrophages within 1% collagen gel, without spheroid.

Percentage of migratory cells (Monocytes, M0, M1 and M2 macrophages) covering the distance (~2 mm) within collagen gel (1%) towards the spheroids. The black line denotes the distance migrated by the cells without the spheroid at bottom.

SI Fig 3

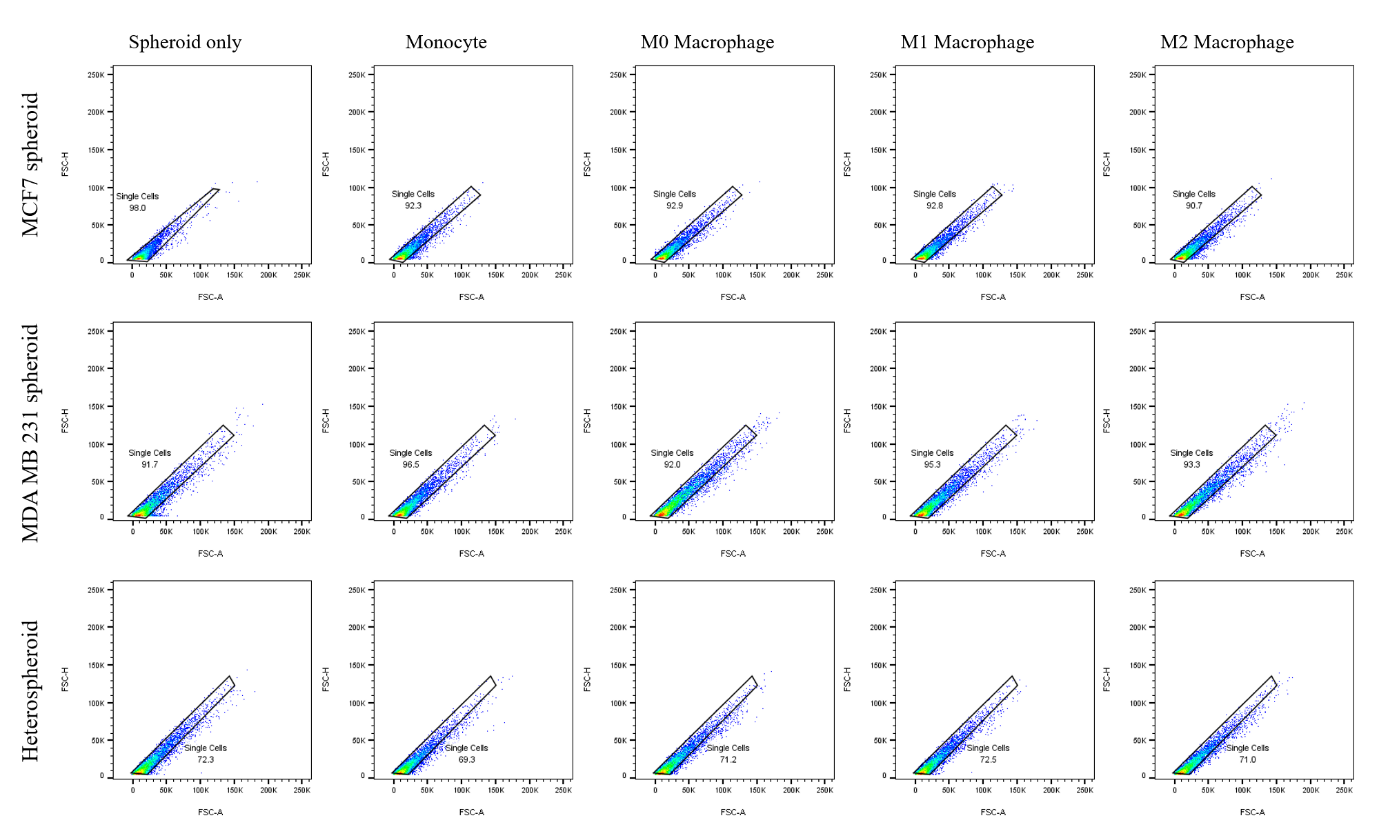

SI Figure 3: Singlet analysis of different cell population (spheroid only and spheroid+ incorporated cells) used for monocyte/macrophage population. The singlet population was further used to separate the green cells (incorporated cells of monocyte/macrophage types).

SI Fig 4

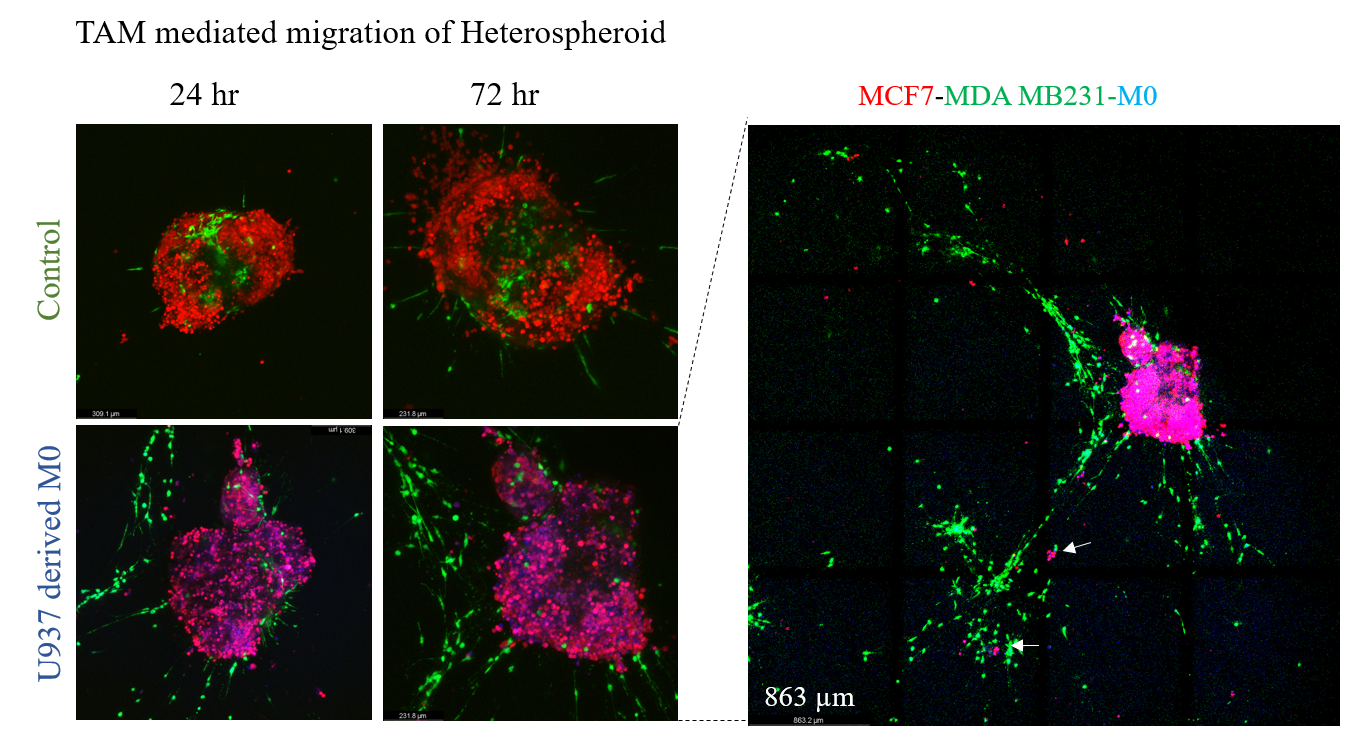

SI Figure 4: Confocal laser microscopic images of control and M0-macrophage incorporated heterospheroids entrapped within 1% collagen for 72 hrs. Very few green cells (MDA MB231) are observed to sprout from the control spheroids after 72 hrs, while in M0-incorporated samples, they are observed to migrate more than 5 mm (well diameter of 96 well plate is 6 mm) as imaged using the Tile imaging option in Leica TC SP8. Red cell clusters (MCF7) are marked using white arrow heads. Scale bar in left panel is 100 µm. Scale bar in right panel is 863 µm.

SI Fig 5

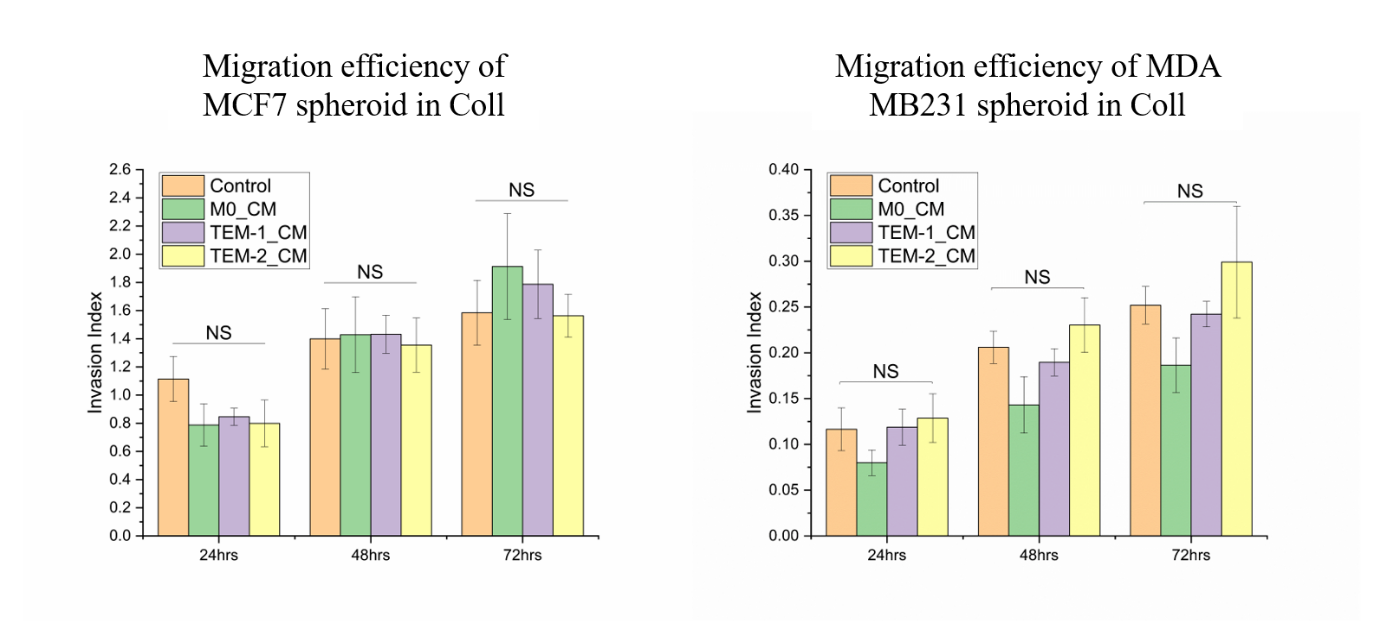

SI Figure 5: Migration efficiency of spheroids (MCF7 and MDA MB231) within the Collagen gel (1%) under the influence of TEM (tumor educated macrophage) mediated conditioned media for 24-72 hr. TEM1 is MCF7 spheroid-CM treated macrophage and TEM2- is MDA MB231 spheroid-CM treated macrophages. No significant difference is observed between the control and test (M0, TEM1, TEM2) samples.

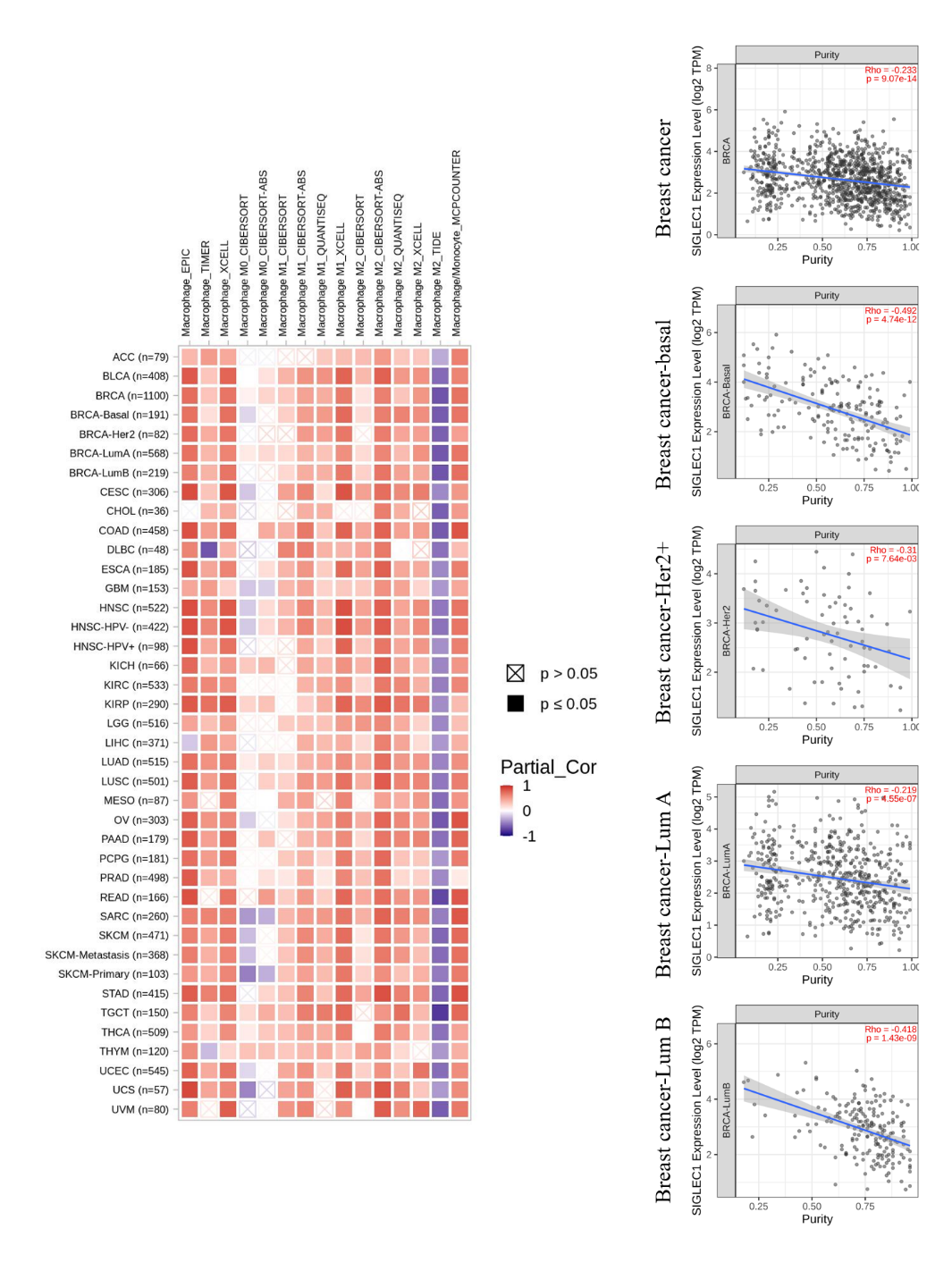
SI Fig 6

SI Figure 6: Correlation study of Siglec-1 expression and macrophage infiltration profile in different types of cancer was done using different algorithm (EPIC, TIMER for example). The correlation type is mentioned by heat map (red-Positive, white-no, and blue-negative correlation). Purity plots of Siglec-1 expression in different breast cancer types was done using TIMER algorithm. Siglec-1 expression is negatively correlated with tumor purity (percentage of malignant cells in a tumor tissue) for each cancer type.

SI Fig 7

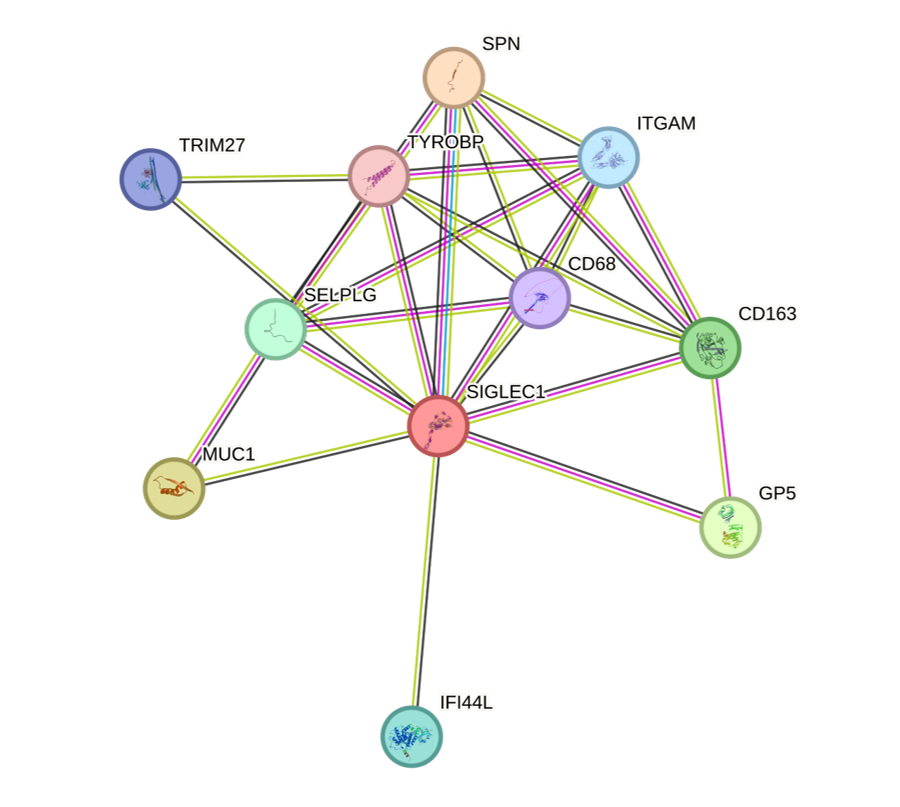

SI Figure 7: STRING analysis of Siglec-1 and its probable binding partners. Siglec-1 is reportedly interact with SPN, MUC1, GP5, CD163, SELPLG, IFI44L, ITGAM, TRIM27, CD68, TYROBP.

SI Fig 8

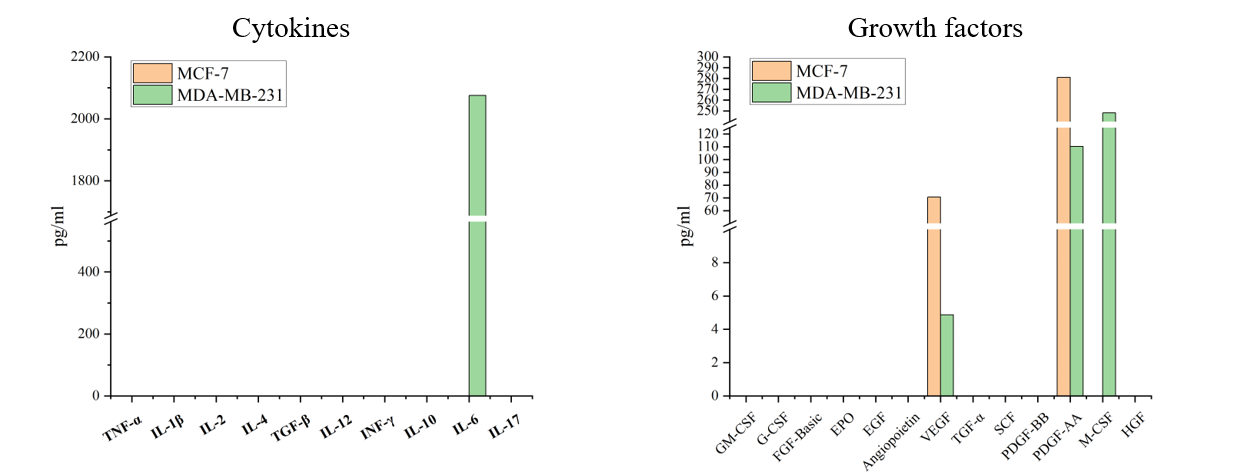

SI Fig 8: Expression level of growth factors and Cytokines were evaluated using spheroid conditioned media through LEGENDplexTM Hu Growth factor Panel and LEGENDplex™ HU Essential Immune Response Panel Cytokine Bead array (SI Table 2 and 3).
